# Control of mitophagy initiation and progression by the TBK1 adaptors NAP1 and SINTBAD

**DOI:** 10.1101/2023.09.25.559255

**Authors:** Elias Adriaenssens, Thanh Ngoc Nguyen, Justyna Sawa-Makarska, Grace Khuu, Martina Schuschnig, Stephen Shoebridge, Emily Maria Watts, Kitti Dora Csalyi, Benjamin Scott Padman, Michael Lazarou, Sascha Martens

## Abstract

Mitophagy preserves overall mitochondrial fitness by selectively targeting damaged mitochondria for degradation. The regulatory mechanisms that prevent PINK1/Parkin-dependent mitophagy and other selective autophagy pathways from overreacting while ensuring swift progression once initiated are largely elusive. Here, we demonstrate how the TBK1 adaptors NAP1 and SINTBAD restrict the initiation of OPTN-driven mitophagy by competing with OPTN for TBK1. Conversely, they promote the progression of NDP52-driven mitophagy by recruiting TBK1 to NDP52 and stabilizing its interaction with FIP200. Notably, OPTN emerges as the primary recruiter of TBK1 during mitophagy initiation, which in return boosts NDP52-mediated mitophagy. Our results thus define NAP1 and SINTBAD as cargo receptor rheostats, elevating the threshold for mitophagy initiation by OPTN while promoting the progression of the pathway once set in motion by supporting NDP52. These findings shed light on the cellular strategy to prevent pathway hyperactivity while still ensuring efficient progression.

## INTRODUCTION

Mitochondria are dynamic and multifunctional organelles, that fuel energy production through oxidative phosphorylation, and play pivotal roles in cell signaling, biosynthetic pathways, and programmed cell death [1–3]. They are susceptible to damage from reactive oxygen species (ROS) and other stressors, necessitating stringent quality control mechanisms [4–6]. The selective removal of damaged mitochondria through autophagy is termed mitophagy and has emerged as essential for maintaining a healthy mitochondrial network [7–12]. Impaired mitophagy links to diverse human disorders, including neurodegenerative diseases, cancer, metabolic syndromes, and aging [13].

The PTEN-induced putative kinase 1 (PINK1) and the E3 ubiquitin ligase Parkin are key players in mitophagy [14, 15], and mutations in these genes underlie early-onset Parkinson’s disease [16–18]. Under basal conditions, PINK1 is continuously degraded by the proteasome [19–21]. However, upon mitochondrial damage, PINK1 accumulates at the outer mitochondrial membrane, recruiting and activating Parkin [22–29]. Parkin marks damaged mitochondria with ubiquitin for recognition by the cargo receptors (also known as cargo adaptors) Optineurin (OPTN) and Nuclear Dot Protein 52 (NDP52, also called CALCOCO2) [30–38]. Autophagosome formation is initiated by the cargo receptors, directly on the surface of the cargo, leading to the engulfment and degradation of the damaged organelle.

TBK1 is a master kinase in mitophagy and other selective autophagy pathways, phosphorylating cargo receptors such as OPTN and NDP52 to increase their affinities for ubiquitin and LC3/GABARAP proteins [32, 33]. However, OPTN and NDP52 utilize TBK1 in different ways. While TBK1 is essential for OPTN-mediated mitophagy initiation [30, 39, 40], NDP52 can redundantly utilize either TBK1 or ULK1 as the mitophagy-initiating kinase [40]. These mechanistic differences between OPTN and NDP52 suggest that TBK1 regulatory factors could play significant roles during mitophagy initiation, especially since OPTN can directly bind TBK1 whereas NDP52 does not [30, 32, 33, 39, 41–44].

NAP1 (also known as AZI2) and SINTBAD are TBK1 adaptors (hereafter referred to as NAP1/SINTBAD), facilitating the interaction between NDP52 and TBK1 in xenophagy—a selective autophagy pathway designed to protect the cytosol against bacterial invasion [42, 45]. NAP1/SINTBAD were found to support NDP52-mediated degradation of *Salmonella enterica* serovar Typhimurium (S. Typhimurium) by interacting with TBK1 and the core autophagy factor FIP200 [42, 43, 46, 47]. However, NAP1/SINTBAD share the same TBK1 binding site as OPTN [44], prompting questions about their potential roles in mitophagy and how their seemingly opposing interactions with OPTN and NDP52 might impact mitophagy dynamics.

We therefore investigated the roles of NAP1/SINTBAD in PINK1/Parkin-dependent mitophagy and discovered their overall inhibitory role in this pathway. While they support NDP52-mediated mitophagy, they negatively regulate TBK1 recruitment and activation by OPTN. This competition for TBK1 binding prevents OPTN from fulfilling one of its primary functions during mitophagy initiation. Our findings highlight a multilayer regulation of mitophagy initiation by NAP1/SINTBAD, acting as cargo receptor rheostats that increase the threshold for mitophagy initiation but promote the progression of the pathway once set in motion. As such, NAP1/SINTBAD provide insight into the cellular strategy that prevents selective autophagy pathways from overreacting while ensuring swift progression once initiated.

## RESULTS

### NAP1/SINTBAD are recruited and co-degraded during mitophagy

To understand whether the TBK1 adaptors NAP1/SINTBAD have a function in PINK1/Parkin mitophagy, we investigated if NAP1/SINTBAD are recruited to mitochondria during this process. To this end, we stably expressed HA-NAP1 or HA-SINTBAD in wild-type (WT) HeLa cells that also expressed YFP-Parkin and assessed their subcellular localization. Under basal conditions, NAP1/SINTBAD were dispersed throughout the cytosol (**Fig. 1A**). However, upon induction of mitophagy using a combination of Oligomycin A and Antimycin A1 (O/A), agents targeting the mitochondrial ATP synthase and complex III, respectively, both NAP1 and SINTBAD notably accumulated on depolarized mitochondria (**Fig. 1A**). We then performed co-staining with WIPI2, a marker for early cup-shaped membrane structures known as phagophores, precursors to autophagosomes. This demonstrated colocalization between NAP1/SINTBAD and WIPI2 (**Fig. 1B**), indicating that both NAP1 and SINTBAD were recruited to sites of autophagosome formation.

**Figure 1.**
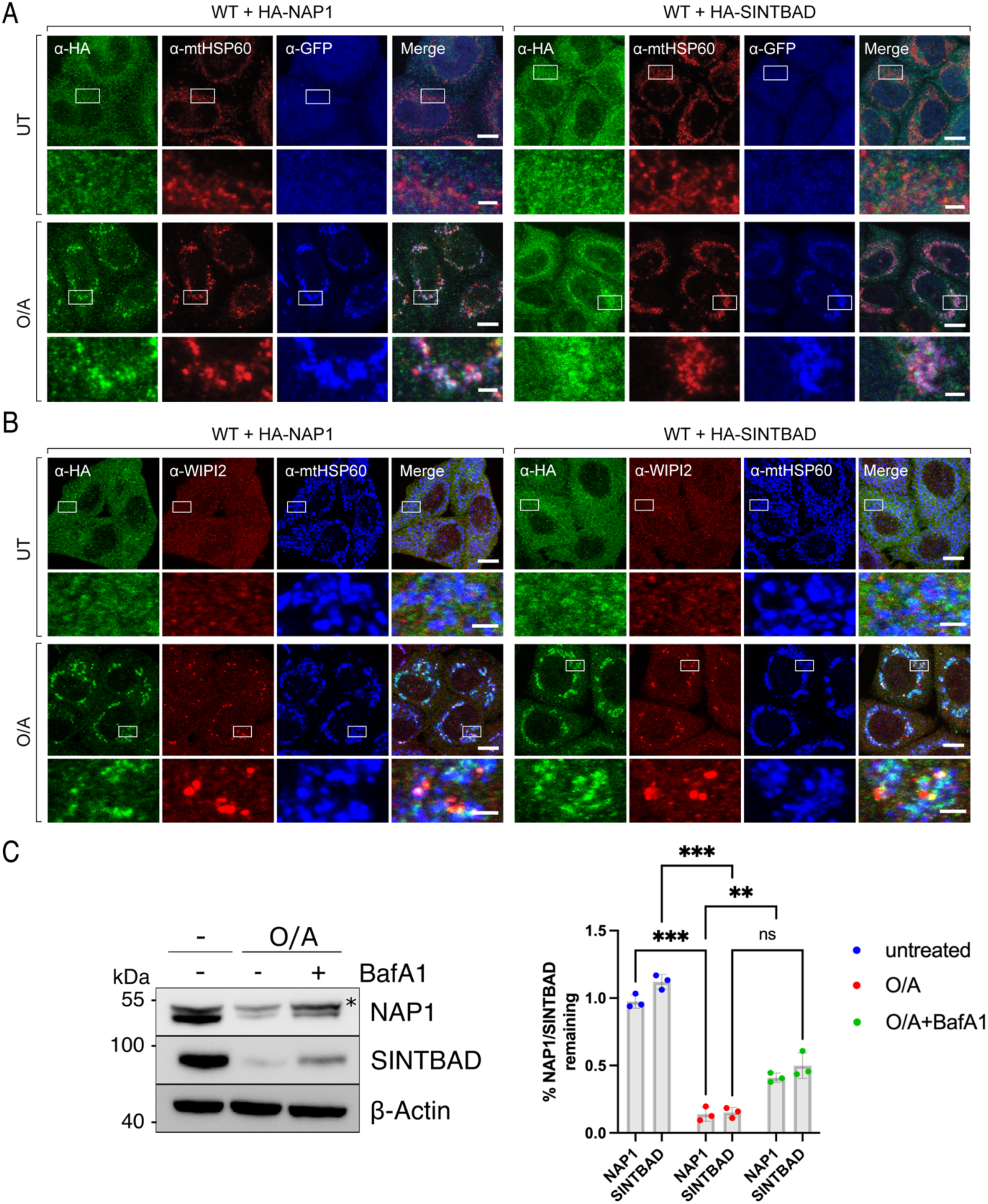
NAP1 and SINTBAD are recruited and co-degraded during mitophagy. (A-B) Wild type (WT) HeLa cells stably expressing YFP-Parkin (A) or BFP-Parkin (B) and HA-NAP1 or HA-SINTBAD, were left untreated or treated with O/A for 2 h, and immunostained with indicated antibodies. (**C**) WT HeLa cells were treated with O/A or O/A and Bafilomycin A1 (BafA1) for 24 h and analyzed by immunoblotting. The levels of NAP1 and SINTBAD were quantified. Asterisks indicates non-specific band. Data in (C) are shown as mean ± s.d. from three independent experiments. Two-way ANOVA with Sidak’s multiple comparison test was performed. \**P*<0.05, \*\**P*<0.005, \*\*\**P*<0.001. ns, not significant. Scale bars: overviews, 10 µm; insets: 2 µm.

To test if NAP1/SINTBAD are degraded along with damaged mitochondria during mitophagy, we assessed the proteins levels of NAP1/SINTBAD. This revealed a decrease in NAP1/SINTBAD levels upon mitophagy induction, which was partially mitigated when lysosomal degradation was inhibited by Bafilomycin A1 (**Fig. 1C**). This indicates that NAP1/SINTBAD are not only recruited to sites of autophagosome formation, but that a portion of NAP1/SINTBAD also undergo autophagy-dependent degradation alongside damaged mitochondria, implying a potential role for them in the PINK1/Parkin mitophagy pathway.

### NAP1/SINTBAD are mitophagy inhibitors

To explore the involvement of NAP1/SINTBAD in PINK1/Parkin mitophagy, we generated knockout HeLa cells for both factors and assessed mitophagy flux. Depletion of either NAP1 or SINTBAD alone did not impact the mitophagy rate in a statistically significant manner, as shown by the mitochondrial-targeted mKeima (mt-mKeima) assay (**Fig. 2A-B**) [48]. Recognizing their structural similarities, which might facilitate compensation for each other, we also generated NAP1/SINTBAD double knockout (DKO) cells. To our surprise, we observed an enhancement in mitophagy flux in NAP1/SINTBAD DKO cells (**Fig. 2C**), contrasting their supporting role in NDP52-mediated xenophagy [47]. This finding was validated by assessing mitochondrial protein COXII levels via western blotting, confirming accelerated mitochondrial degradation in NAP1/SINTBAD DKO cells (**Fig. 2D**).

**Figure 2.**
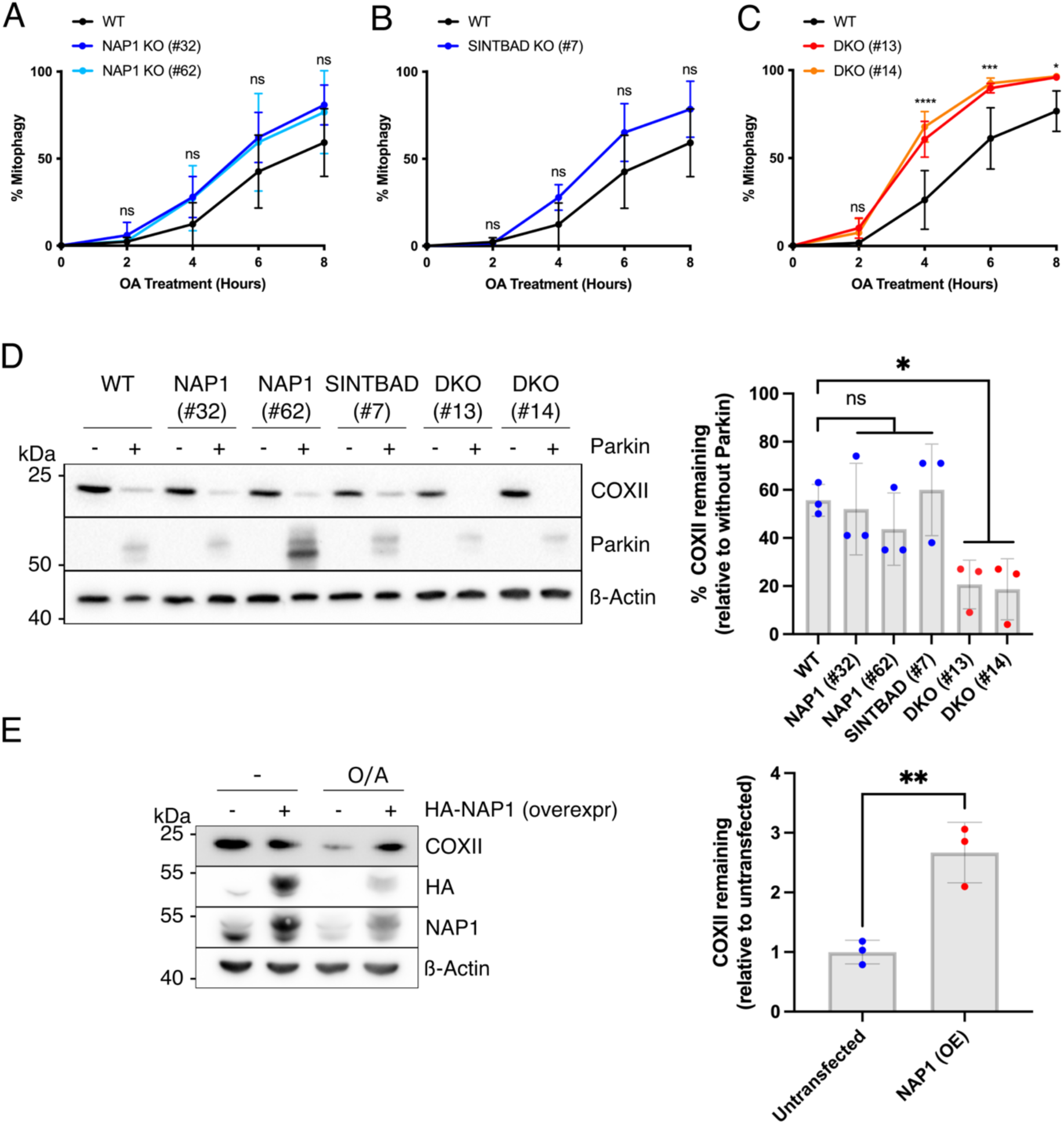
NAP1 and SINTBAD are negative regulators of mitophagy. (**A-C**) Mitophagy flux was measured by flow cytometry in indicated HeLa cell lines expressing YFP-Parkin and mt-mKeima, untreated or treated with O/A for indicated times; wild-type versus NAP1 KO (A), SINTBAD KO (B), or NAP1 and SINTBAD double knockout (DKO) cells (C). (**D**) Immunoblotting of COXII levels in various HeLa cell lines treated with O/A for 18 h. PINK1/Parkin-dependent versus PINK1/Parkin-independent mitophagy was compared by overexpression of YFP-Parkin. The percentage of COXII remaining was quantified. (**E**) Immunoblotting of COXII levels in HeLa cells overexpressing (OE) HA-NAP1 and treated with O/A for 16 h. The proportion of COXII remaining after O/A relative to the untransfected sample was quantified. Data in (A-E) are shown as mean ± s.d. from three independent experiments. Two-way analysis of variance [ANOVA] (A-C); One-way ANOVA with Dunnett’s multiple comparison test was performed in (D) and a two-tailed unpaired Student’s *t* test in (E). \**P*<0.05, \*\**P*<0.005, \*\*\**P*<0.001, \*\*\*\**P*<0.0001. ns, not significant.

To substantiate their inhibitory role, we investigated whether NAP1 overexpression could inhibit mitophagy. Our analysis indeed revealed that NAP1 overexpression led to reduced COXII degradation (**Fig. 2E**). Thus, NAP1/SINTBAD serve as mitophagy inhibitors, counteracting PINK1/Parkin-mediated mitophagy.

To explore whether NAP1/SINTBAD also regulate non-selective bulk autophagy, we evaluated p62 degradation in starved cells. Our findings indicated no discernible changes in p62 degradation in single or double knockout cell lines when compared to control wild-type cells (**Fig. S1**). Therefore, NAP1/SINTBAD are involved in the regulation of selective forms of autophagy, such as mitophagy, but not in non-selective bulk autophagy.

### NAP1/SINTBAD support NDP52-mediated mitophagy by enabling TBK1 binding and stabilizing interactions with the autophagy machinery

To investigate the mechanisms underlying NAP1/SINTBAD’s inhibition of mitophagy, we first focused on their functional interaction with NDP52, as they were previously implicated in an NDP52-dependent selective autophagy pathway, albeit in a stimulatory manner [47].

To explore their interplay with NDP52, we generated CRISPR/Cas9 double knockout clones for NAP1/SINTBAD in the pentaKO background, which lacks five key cargo receptors OPTN, NDP52, TAX1BP1, p62, and NBR1 [30]. This allowed us to reintroduce NDP52 into these cells and to assess NDP52-driven mitophagy rates in the presence or absence of NAP1/SINTBAD, eliminating the confounding effects from other cargo receptors, including OPTN. Surprisingly, contrary to our previous observations (**Fig. 2**), deleting NAP1/SINTBAD in these cells resulted in reduced mitophagy. This was evident from reduced degradation of the mitochondrial marker COXII (**Fig. 3A**), decreased mt-mKeima conversion (**Fig. 3B**), and impaired TBK1 activation (**Fig. 3C**). Moreover, the deletion of NAP1/SINTBAD may have weakened the NDP52-FIP200 interaction, as *in vitro* reconstitution of NDP52-mediated mitophagy initiation revealed that SINTBAD enhanced the NDP52-FIP200 interaction (**Fig. 3D**), underscoring an important role for NAP1/SINTBAD in this critical early step of mitophagy initiation.

**Figure 3.**
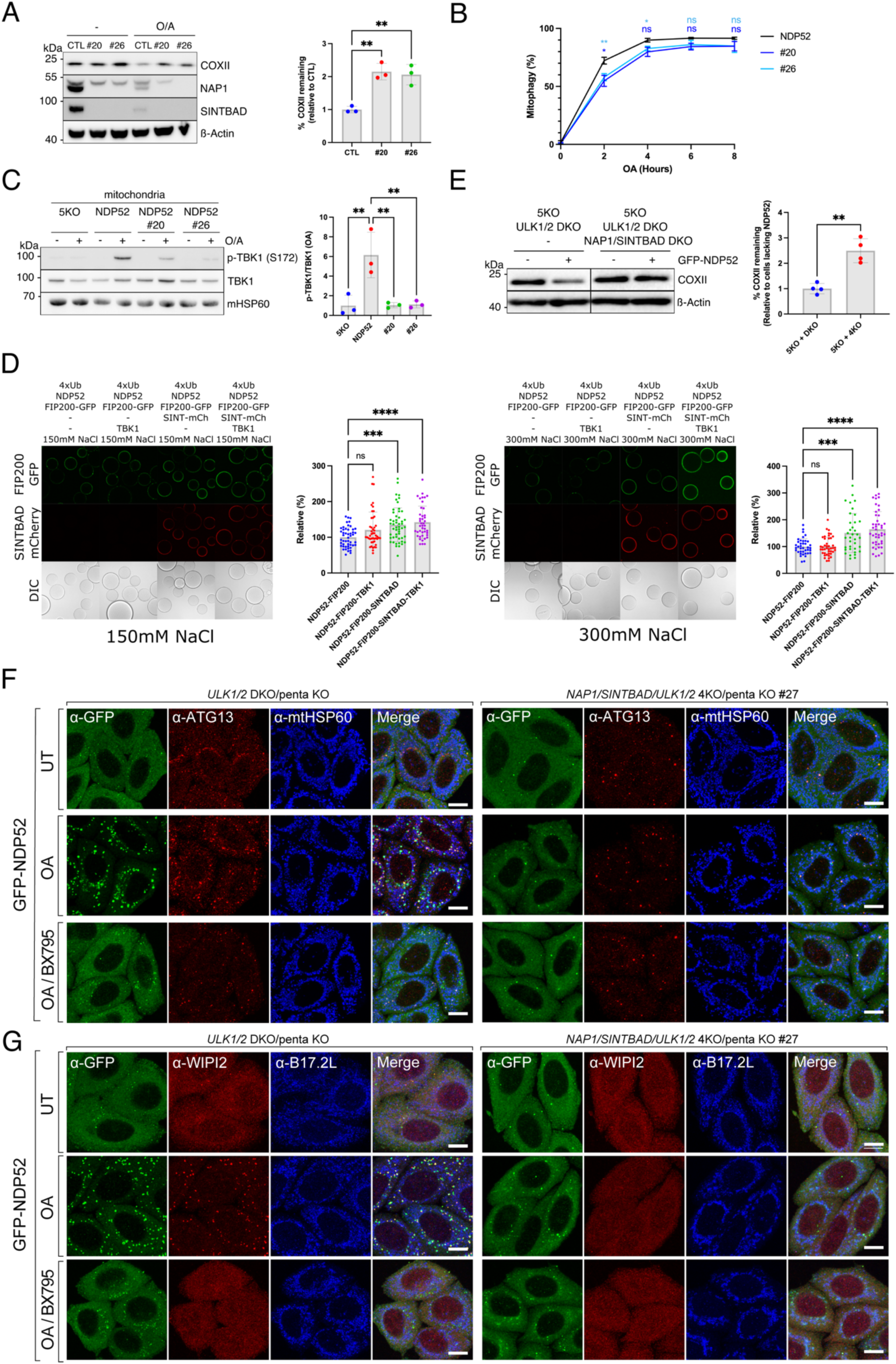
NAP1 and SINTBAD support NDP52-mediated mitophagy by stabilizing interactions with autophagic machinery. (**A**) Penta KO (parental control, CTL) and NAP1/SINTBAD DKO/penta KO (clones 20 and 26) expressing BFP-Parkin and GFP-NDP52 were treated with O/A for 16 h and analyzed by immunoblotting. The percentage of COXII remaining was quantified. (**B**) Indicated cell lines expressing BFP-Parkin and mt-mKeima, were treated with O/A for indicated times. Mitochondrial flux was measured by flow cytometry. Representative FACS plots are provided in Figure S2. (**C**) Crude mitochondria were isolated from penta KO and NAP1/SINTBAD DKO/penta KO (clones 20 and 26) expressing BFP-Parkin and GFP-NDP52 untreated or treated with O/A for 1 h and analyzed via immunoblotting with indicated antibodies. The fraction of p-TBK1 over total TBK1 was quantified. (**D**) Biochemical reconstitution of mitophagy initiation by NDP52. Glutathione Sepharose beads coated with GST-tagged linear ubiquitin chains (GST-4×Ub) were incubated with NDP52, SINTBAD-mCherry, TBK1, and FIP200-GFP, as indicated, in bead assay buffer containing either 150 mM or 300 mM NaCl and supplemented with ATP/MgCl_2_. Samples were analyzed by confocal imaging. (**E**) Penta KO with ULK1/2 DKO and penta KO with ULK1/2/NAP1/SINTBAD 4KO (clones 13 and 27) expressing BFP-Parkin and GFP-NDP52 were treated with O/A for 16 h and analyzed by immunoblotting. The percentage of COXII remaining was quantified. (**F-G**) Penta KO with ULK1/2 DKO and penta KO with ULK1/2/NAP1/SINTBAD 4KO HeLa cells stably expressing BFP-Parkin were left untreated or treated with O/A or O/A plus TBK1 inhibitor (BX795) for 1 h, and immunostained with indicated antibodies. Note the defect in GFP-NDP52 recruitment in #27 due to failure of recruiting downstream ATG8-molecules, which feedback and stabilize NDP52 (Padman et al. 2019). Data in (A-E) are shown as mean ± s.d. from three independent experiments. Each data point in (E) represents the mean signal intensity for an individual bead. One-way ANOVA with Dunnett’s multiple comparison test was performed in (A, C, and D), and two-way ANOVA with Turkey’s multiple comparisons test in (B). \**P*<0.05, \*\**P*<0.005, \*\*\**P*<0.001, \*\*\*\**P*<0.0001. ns, not significant.

Despite the significant contribution of NAP1/SINTBAD to these important first steps of NDP52-mediated mitophagy initiation, the overall reduction in mitophagy flux was relatively modest. However, considering that NDP52 can drive mitophagy through either ULK1/2 or TBK1 [40], we knocked out ULK1/2 in NAP1/SINTBAD DKO/pentaKO cells to elucidate the necessity of NAP1/SINTBAD when NDP52 engages in mitophagy solely through the TBK1 pathway. In the absence of ULK1/2, NAP1/SINTBAD emerged as essential factors for NDP52-mediated mitophagy, evident from significantly reduced COXII turnover (**Fig. 3E**) and diminished WIPI2 and ATG13 recruitment upon O/A treatment (**Fig. 3F-G**). The latter to the same extent as when we inhibited TBK1 with the small molecule BX795.

In summary, these findings reveal important roles for NAP1/SINTBAD in supporting NDP52-mediated mitophagy through the recruitment of TBK1 and stabilization of the NDP52-FIP200 complex.

### NAP1/SINTBAD are sufficient to induce mitophagy when recruited to mitochondria

From our experiments above (**Fig. 3**), it becomes evident that NAP1/SINTBAD exhibit traits of cargo receptors, including their ability to bind FIP200 and TBK1, albeit lacking the ubiquitin binding capabilities of cargo receptors. However, ubiquitin chains are critical in marking damaged organelles for autophagic degradation. With this in mind, we hypothesized that bypassing this ubiquitin-dependent recruitment by artificially tethering NAP1 to the outer mitochondrial membrane might be sufficient to initiate autophagosome biogenesis.

To test this hypothesis, we employed a chemically induced dimerization (CID) assay, wherein FRB and FKBP can be dimerized upon rapalog addition [49, 50]. By positioning FRB on the mitochondrial outer membrane through fusion with the transmembrane domain of FIS1 and attaching NDP52 or NAP1 to FKBP, we gained the ability to redirect NAP1 or NDP52 to the outer mitochondrial membrane upon rapalog treatment (**Fig. 4A**).

**Figure 4.**
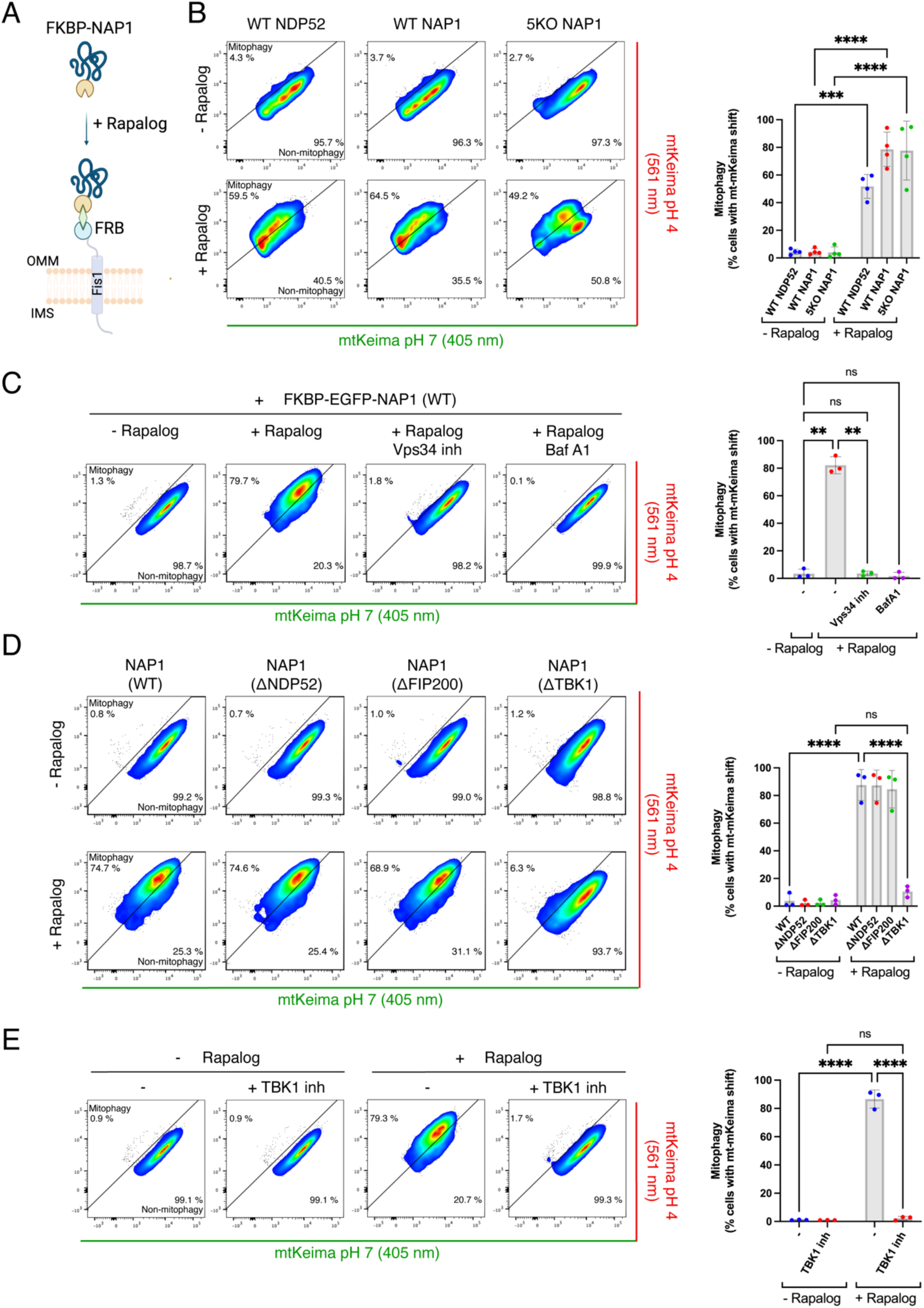
NAP1 can drive mitophagy when artificially tethered to the mitochondrial surface. (**A**) Diagram of the experimental set-up and the effect of rapalog treatment, resulting in the tethering of NDP52 or NAP1 to the outer mitochondrial membrane. IMS: intermembrane space, OMM: outer mitochondrial membrane. (**B**) Mitophagy flux was measured by flow cytometry in wild-type (WT) or penta KO (5KO) HeLa cells expressing BFP-Parkin and mt-mKeima, not induced or induced for 24 h by rapalog treatment. (**C**) As in (B) but with and without the addition of autophagy inhibitors: PI3K inhibitor (Vps34 inh) and Bafilomycin A1 (Baf A1). (**D**) Different NAP1 variants deficient in binding NDP52, FIP200, or TBK1 were ectopically tethered to the outer mitochondrial membrane, and the level of mitophagy induction was compared to wild-type NAP1. (**E**) As in (B), with the pentaKO background, but with and without the addition of the TBK1 inhibitor (GSK8612). Representative FACS plots are shown from one of four (B) or three (C-E) replicates. The percentage of non-induced cells (lower right) versus mitophagy-induced cells (upper left) is indicated. Two-way ANOVA with Turkey’s multiple comparisons test in (B,C,D,E). \**P*<0.05, \*\**P*<0.005, \*\*\**P*<0.001, \*\*\*\**P*<0.0001. ns, not significant.

We first confirmed that NDP52 induced mitophagy upon rapalog addition (**Fig. 4B**), as previously demonstrated [30, 51]. We then evaluated whether FKBP-NAP1 could similarly initiate mitophagy. Intriguingly, artificial tethering of NAP1 to the mitochondrial surface resulted in comparable levels of mitophagy induction upon rapalog treatment as compared to NDP52 (**Fig. 4B**). To rule out that this effect stemmed from the indirect recruitment of NDP52 by NAP1, we repeated the experiment in pentaKO cells. This confirmed that NAP1 could autonomously induce mitophagy, independently of NDP52 (**Fig. 4B**). Moreover, blocking autophagosome formation with a Vps34 inhibitor or impeding autophagosome degradation with Bafilomycin A1 validated that the mitochondrial turnover was mediated by autophagy (**Fig. 4C**).

Using the rapalog-induced tethering assay, we further dissected the mechanism of NAP1-induced mitophagy. Specifically, we utilized NAP1 mutants deficient in NDP52-binding, FIP200-binding, or TBK1-binding. The mutants lacking NDP52-or FIP200-binding abilities retained their capacity to induce mitophagy upon rapalog treatment (**Fig. 4D**). However, the TBK1-binding deficient mutant lost its ability to initiate mitophagy, underscoring the critical role of the NAP1-TBK1 interaction in mitophagy. Consistently, inhibition of TBK1 with the small molecule GSK8612 prevented ectopically tethered NAP1 from inducing mitophagy (**Fig. 4E**). Collectively, these findings highlight the resemblance of NAP1/SINTBAD to cargo receptors, with the exception of ubiquitin binding. By artificially tethering NAP1 to the mitochondrial surface, we demonstrated its competency as an autophagy cargo receptor in a TBK1-dependent manner. Based on these insights, we propose the term “cargo co-receptors” for NAP1/SINTBAD, emphasizing their ability to facilitate selective autophagy through interactions with cargo receptors like NDP52.

### NAP1/SINTBAD restrict mitophagy by competing with OPTN for TBK1 binding and activation

While our findings above underscore the importance of NAP1/SINTBAD for NDP52-driven selective autophagy pathways, these results do not explain our earlier observations in NAP1/SINTBAD DKO cells, where their overall effect on mitophagy was inhibitory rather than stimulatory. This suggests that the roles of NAP1/SINTBAD in mitophagy might be cargo receptor-specific, considering that NAP1/SINTBAD DKO cells express all five cargo receptors, while experiments in the pentaKO background were conducted in cells expressing only NDP52. Based on the fact that NAP1/SINTBAD bind to TBK1 at the same binding site as OPTN [44], we hypothesized that their inhibitory impact on mitophagy might arise from direct or indirect regulation of OPTN, the other major cargo receptor in PINK1/Parkin-dependent mitophagy.

To test whether NAP1/SINTBAD could inhibit mitophagy by competing with OPTN for TBK1 binding, we reconstituted the initiation of OPTN-driven mitophagy *in vitro* using purified components. Agarose beads coated with linear 4× ubiquitin, mimicking the surface of ubiquitin-marked damaged mitochondria, were co-incubated with mCherry-tagged OPTN, EGFP-tagged TBK1, and increasing concentrations of NAP1 (**Fig. 5A**). This experiment revealed that OPTN was recruited to the ubiquitin-coated beads, subsequently recruiting TBK1 (**Fig. 5B**). However, increasing NAP1 levels led to TBK1 displacement from the OPTN-bound beads, indicating that OPTN and NAP1 compete for the same binding site. This competition was further validated through conventional pull-down experiments (**Fig. S3**).

**Figure 5.**
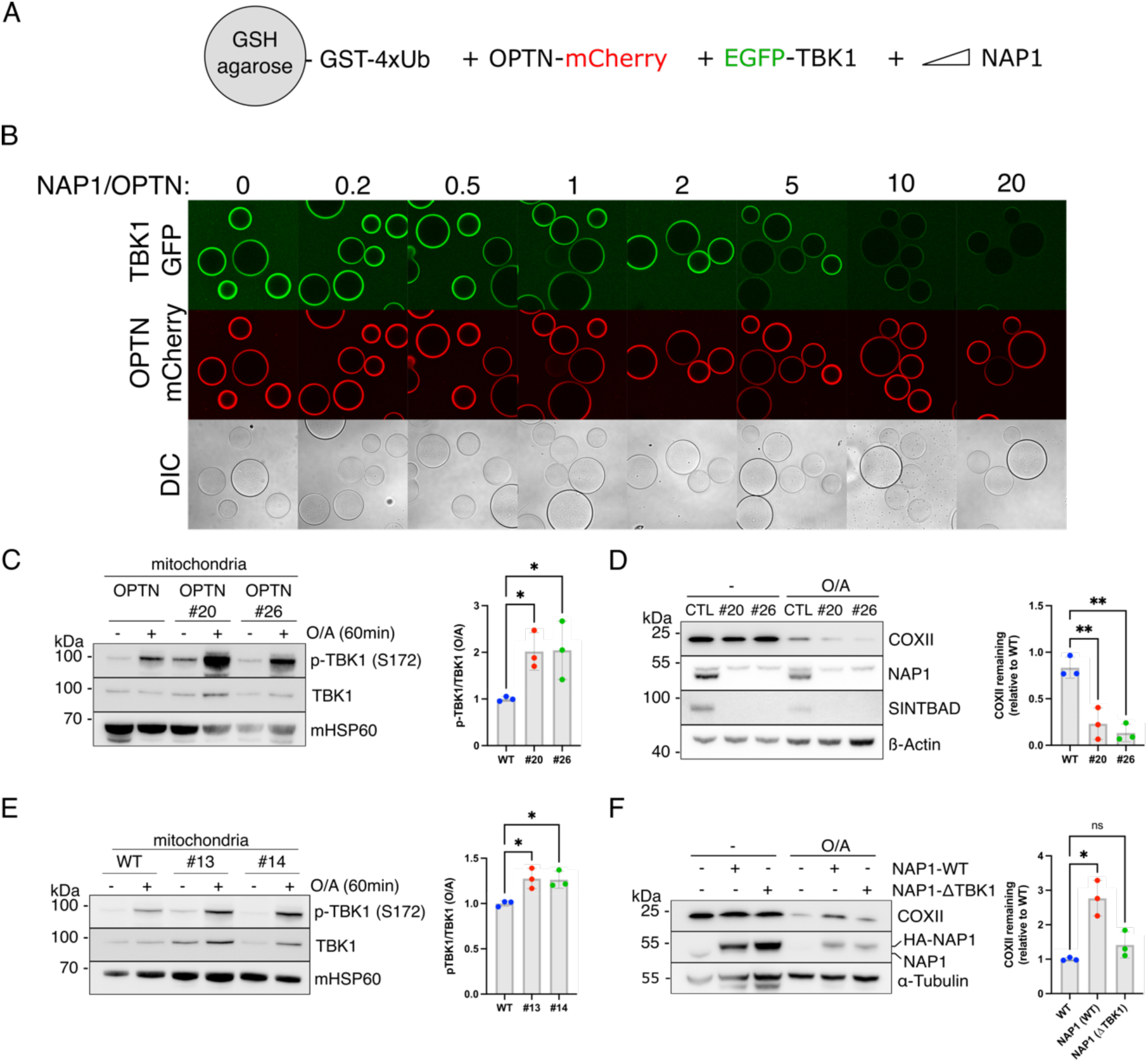
NAP1 and SINTBAD compete with OPTN for TBK1 binding. (**A**) Diagram of the experimental set-up to assess competition between OPTN and NAP1 for TBK1 binding. (**B**) Biochemical reconstitution of the recruitment of GFP-TBK1 by mCherry-OPTN to GST-4×Ub coated beads in the absence or presence of increasing amounts of NAP1. The experiment was performed without ATP, and samples were analyzed by confocal imaging. One of three representative experiments is shown. (**C**) Crude mitochondria were isolated from penta KO and NAP1/SINTBAD DKO/penta KO (clones 20 and 26) expressing BFP-Parkin and GFP-OPTN untreated or treated with O/A for 1 h and analyzed via immunoblotting with indicated antibodies. The fraction of p-TBK1 over total TBK1 was quantified. (**D**) Immunoblotting of COXII levels in Penta KO (parental control, CTL) and NAP1/SINTBAD DKO/penta KO (clones 20 and 26) expressing BFP-Parkin and GFP-OPTN, and treated with O/A for 16 h, analyzed by immunoblotting. The percentage of COXII remaining was quantified. The upper band for NAP1 is non-specific. (**E**) Crude mitochondria were isolated from wild-type (WT) HeLa cells and NAP1/SINTBAD DKO cells in WT background (clones #13 and #14) expressing BFP-Parkin untreated or treated with O/A for 60 min were analyzed by immunoblotting with indicated antibodies. The fraction of p-TBK1 over total TBK1 was quantified. (**F**) Wild-type HeLa cells expressing BFP-Parkin were stably transduced with HA-NAP1 wild-type (WT) or TBK1-binding deficient mutant (ΔTBK1). Cells were treated with O/A for 16 h and analyzed by immunoblotting. The percentage of COXII remaining after O/A treatment was quantified. Data are shown as mean ± s.d. from three independent experiments. One-way ANOVA with Dunnett’s multiple comparison test was performed. \**P*<0.05, \*\**P*<0.005.

To assess whether NAP1/SINTBAD also competed with OPTN for TBK1 binding in cells, we used the NAP1/SINTBAD DKOs in the pentaKO background, where OPTN was reintroduced. This setup allowed us to distinguish the effects of NAP1/SINTBAD on OPTN-mediated mitophagy from those on NDP52-mediated mitophagy. Following mitophagy induction in these cells, we observed increased TBK1 activation as indicated by higher levels of p-S172 TBK1 in the absence of NAP1/SINTBAD (**Fig. 5C**). This suggests that NAP1/SINTBAD suppress TBK1 activation in OPTN-mediated mitophagy, consistent with their competition with OPTN for TBK1 binding (**Fig. 5B**). We then examined whether increased TBK1 activation resulted in accelerated mitophagy. Indeed, measurements of mitophagy levels, indicated by COXII degradation in pentaKO cells rescued with OPTN, confirmed the acceleration of OPTN-driven mitophagy in the absence of NAP1/SINTBAD (**Fig. 5D**). Consistently, we also detected accelerated mitophagy using the mt-mKeima assay (**Fig. S4**).

Next, we quantified the amount of activated TBK1 relative to total TBK1 on the surface of purified mitochondria in wild-type cells versus NAP1/SINTBAD DKO cells expressing all five cargo receptors. This revealed increased TBK1 activation upon NAP1/SINTBAD deletion (**Fig. 5E**), suggesting that NAP1/SINTBAD are indeed competing with OPTN for TBK1 binding in the cell.

To further validate that NAP1/SINTBAD inhibit mitophagy, at least in part, through competition for TBK1 binding, we engineered a NAP1 mutant (L226Q/L233Q) deficient in TBK1 binding (**Fig. S5**) and assessed its inhibitory potential. Upon overexpression of wild-type NAP1 or the TBK1-binding deficient mutant in wild-type HeLa cells, we observed that wild-type NAP1 reduced the overall COXII degradation, as observed earlier (**Fig. 2E**). However, this effect was nearly completely abolished for the TBK1-binding deficient mutant (**Fig. 5F**), further supporting the notion that NAP1/SINTBAD restrict mitophagy initiation through competition for TBK1 binding.

Taken together, these findings reveal that NAP1/SINTBAD, through competition for TBK1 binding, can restrict the initiation of mitophagy.

### OPTN is the primary recruiter and activator of TBK1 during mitophagy initiation

The insights gathered above not only unveil a novel regulatory step at the onset of mitophagy, but also shed light on a critical role for TBK1 in ensuring the efficient progression of mitophagy. Our results show that NAP1/SINTBAD restrict mitophagy initiation by limiting the TBK1 recruitment by OPTN, hinting at a dominant role for OPTN in recruiting and activating TBK1. We therefore set out to dissect the underlying mechanisms of TBK1 recruitment and activation during mitophagy.

Consistent with prior research, we first confirmed that the activation of TBK1 strictly relies on the presence of cargo receptors, as their absence resulted in the absence of TBK1 activation (**Fig. 6A**) [30]. Furthermore, in line with the mechanism by which TBK1 is activated through local clustering on the ER surface by the cGAS-STING complex [52–56], we find that TBK1 is also activated locally on the mitochondrial surface during mitophagy (**Fig. 6B**). This aligns with the requirement of TBK1 dimers to be brought into close proximity, enabling trans autophosphorylation, as the kinase domain cannot access the activation loop in *cis* and the two kinase domains in the dimer face away from one another [57–59]. Our data thus propose an essential role for cargo receptors in locally clustering TBK1 dimers on the mitochondrial surface. This is consistent with a recently proposed model, positing that TBK1 is activated from a local platform of OPTN molecules [60].

**Figure 6.**
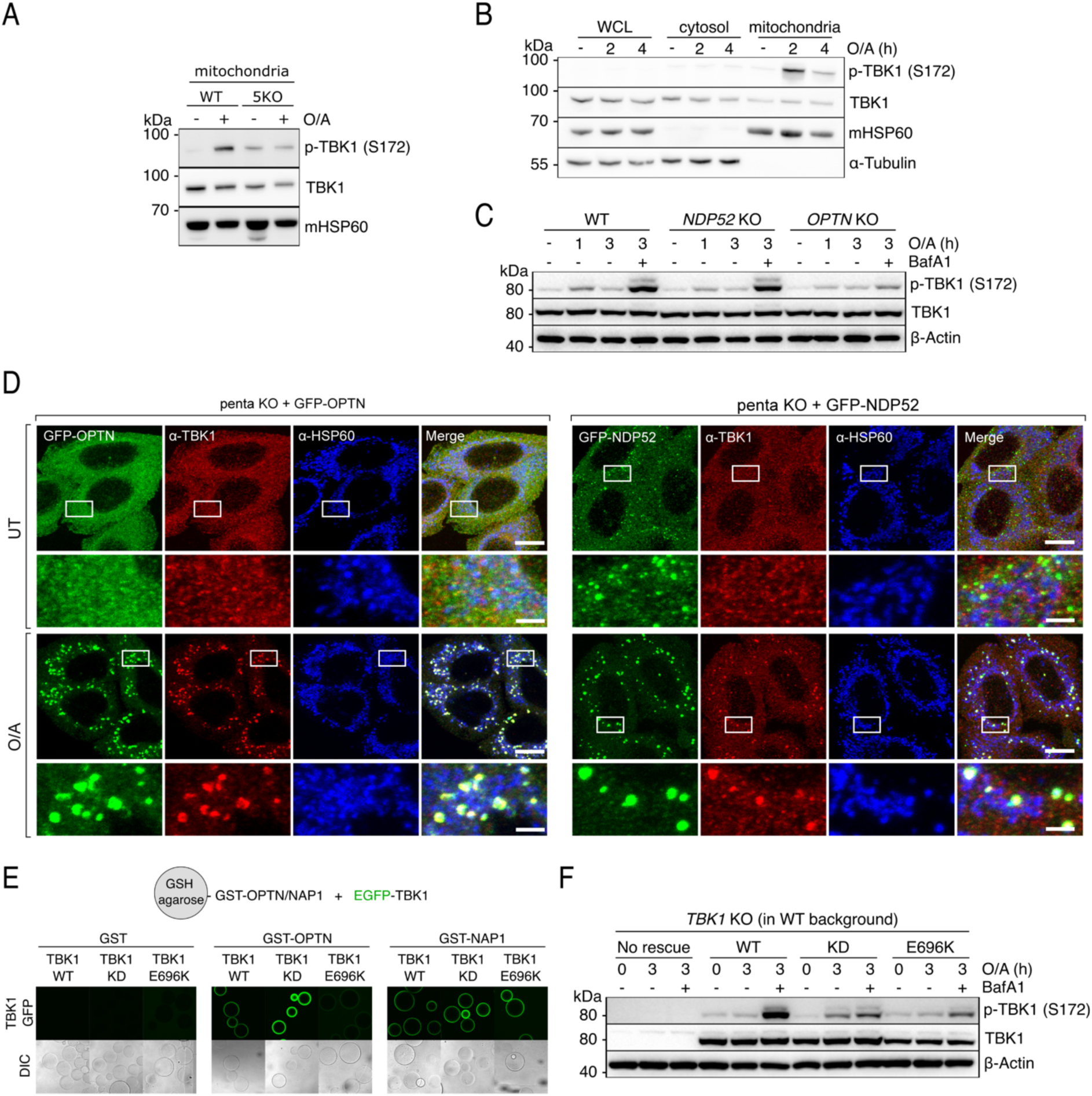
OPTN is the primary recruiter and activator of TBK1 during mitophagy initiation. (**A**) Immunoblotting of crude mitochondrial fraction isolated from wild-type versus penta KO (5KO) cells, expressing BFP-Parkin and treated with O/A for 6 h. (**B**) Crude mitochondria were isolated from wild-type HeLa cells expressing BFP-Parkin, treated with O/A for the indicated times, and compared to the cytosolic and whole cell lysate (WCL) fractions via immunoblotting with indicated antibodies. (**C**) Immunoblotting of phosphorylated TBK1 (S172) in wild-type versus OPTN or NDP52 knockout cells expressing BFP-Parkin treated with O/A in the absence or presence of Bafilomycin A1 (BafA1) for the indicated times and analyzed by immunoblotting. (**D**) Penta KO HeLa cells stably expressing BFP-Parkin and rescued with GFP-OPTN or GFP-NDP52 were either left untreated or treated with O/A for one hour and immunostained with indicated antibodies. (**E**) In vitro binding assay using glutathione-coupled agarose beads coated with GST, GST-OPTN, or GST-NAP1 and incubated with EGFP-TBK1 wild-type (WT), EGFP-TBK1 kinase-dead (KD), or EGFP-TBK1 E696K mutant (E696K). Samples were analyzed by confocal microscopy. (**F**) Immunoblotting of TBK1 knockout HeLa cells that were either not rescued, rescued with TBK1 wild-type (WT), TBK1 kinase-dead (KD), or TBK1 E696K mutant (E696K) and treated with O/A in the absence or presence of Bafilomycin A1 (BafA1) for the indicated times. Data are shown as representative of one of three replicates.

To test whether TBK1 activation predominantly relies on OPTN, as implied by our NAP1/SINTBAD results, we compared TBK1 activation in wild-type HeLa cells to cells lacking either OPTN or NDP52. This comparison revealed a severe reduction in TBK1 activation upon OPTN deletion as evident from decreased TBK1 phosphorylation (**Fig. 6C**). In contrast, NDP52 deletion had a relatively minor impact on TBK1 activation (**Fig. 6C**). We then compared the amount of TBK1 recruitment during OPTN-versus NDP52-driven mitophagy by rescuing the pentaKO cells with either OPTN or NDP52. This revealed OPTN’s pronounced ability to recruit TBK1 to the mitochondrial surface upon mitophagy induction, while NDP52 recruited TBK1 to a lesser extent (**Fig. 6D**).

To further corroborate this result, we employed the ALS-causing TBK1 E696K mutation. This mutant failed to bind OPTN *in vitro*, in line with prior research [39, 44, 61, 62]. However, this mutation retained its binding capacity to NAP1 (**Fig. 6E**). In wild-type HeLa cells, expressing both OPTN and NDP52, the TBK1 E696K mutant was previously shown to be no longer recruited to damaged mitochondria [39, 61]. Consistently, we show that this is accompanied by a drastic reduction of TBK1 activation (**Fig. 6F**), reinforcing the importance of clustering for TBK1 activation. Moreover, these findings are also consistent with OPTN playing a primary role in recruiting and clustering TBK1 on the mitochondrial surface, which cannot be sufficiently compensated for by the NDP52-NAP1/SINTBAD axis in HeLa cells. This underscores the importance of OPTN-mediated TBK1 recruitment.

Together, our results provide evidence for a crucial role of OPTN in recruiting and activating TBK1 during mitophagy, explaining how interference with this interaction by NAP1/SINTBAD can effectively restrict mitophagy initiation.

### Crosstalk between the OPTN-axis and NDP52-axis stimulates mitophagy

We wondered whether the crucial role of OPTN in TBK1 activation might also influence the NDP52 axis. Previous research revealed that either cargo receptor alone is sufficient to initiate mitophagy [30]. However, several tissues express both cargo receptors. In tissues such as the brain, where NDP52 expression is low [30], the NDP52-related protein TAX1BP1 is expressed. We therefore hypothesized that a crosstalk might exist between OPTN and NDP52, allowing each receptor to leverage its strengths so that their combined presence results in robust mitophagy control and progression.

To be able to test this, we designed a system that enabled us to exploit OPTN’s capacity to recruit TBK1 during mitophagy, but omitting its ability to interact with other components of the autophagy machinery [41, 63, 64]. To this end, we created a rapalog-induced dimerization assay, linking only the minimal sequence of OPTN (residues 2-119) essential for TBK1 binding to FKBP (**Fig. 7A**). Using this system in the pentaKO background, we tested whether this truncated OPTN fragment could effectively recruit and activate TBK1 at the mitochondrial surface. Indeed, purification of mitochondria from rapalog-treated HeLa cells revealed that rapalog induced the translocation of FKBP-OPTN(2-119) and TBK1 (**Fig. 7B**). Crucially, the recruitment of TBK1 to the mitochondrial surface was sufficient to induce TBK1 activation, as demonstrated by the increase in phosphorylated TBK1 in the mitochondrial fraction upon rapalog treatment.

**Figure 7.**
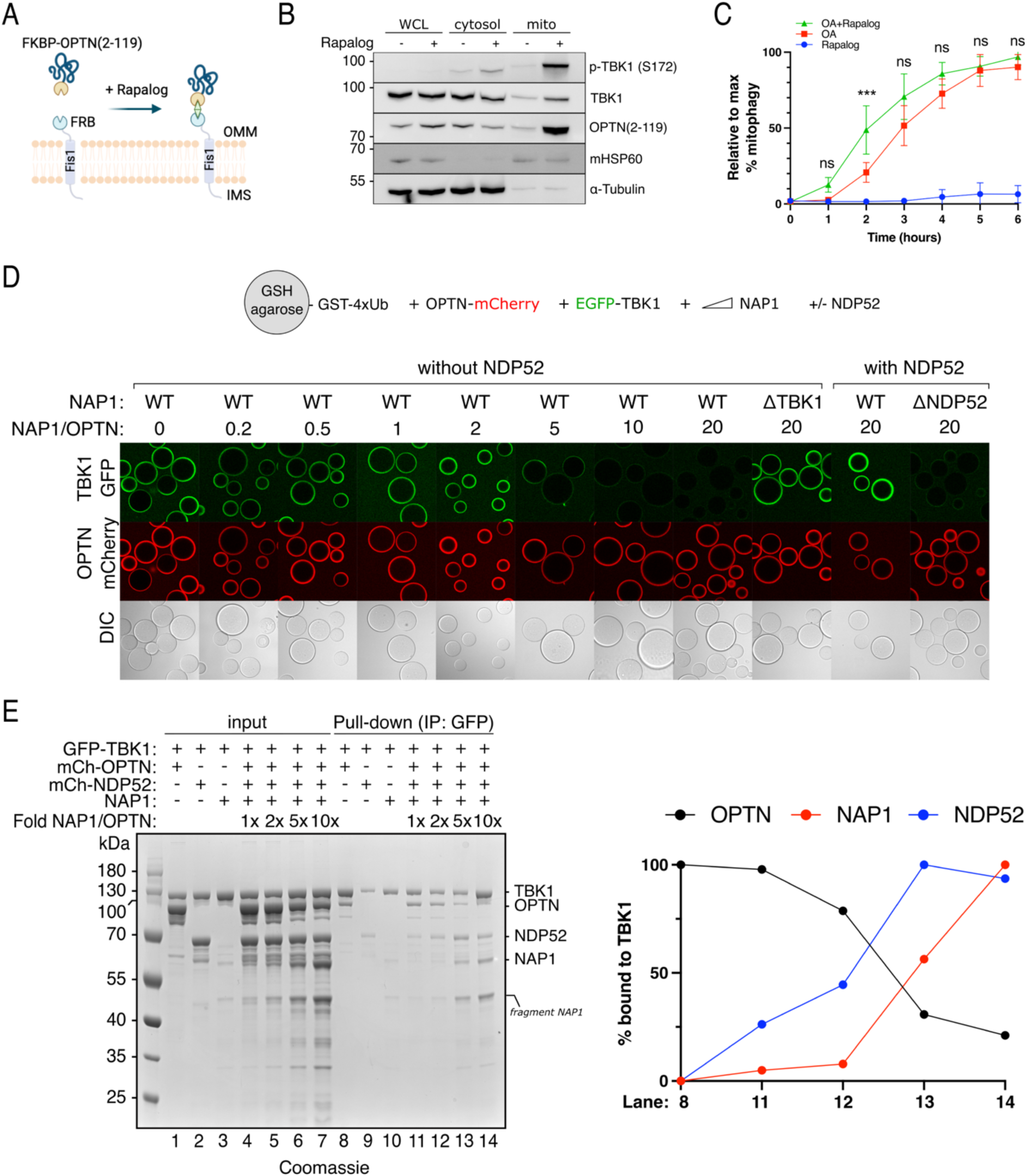
Crosstalk between the OPTN-axis and NDP52-axis stimulates mitophagy. (**A**) Diagram of the experimental set-up and the effect of rapalog treatment, resulting in the tethering of FKBP-OPTN(2-119) to the outer mitochondrial membrane. IMS: intermembrane space, OMM: outer mitochondrial membrane. (**B**) Whole cell lysate (WCL), cytosol, and mitochondrial (mito) fractions from untreated versus 3 h rapalog-treated cells were analyzed by immunoblotting for phosphorylated TBK1 (S172) and other indicated antibodies. (**C**) Penta KO cells expressing BFP-Parkin and GFP-NDP52 were further transduced with Fis1-FRB and FKBP-EGFP-OPTN(2-119), and treated with rapalog alone, O/A alone, or rapalog plus O/A for the indicated times. Mitophagy flux was measured by flow cytometry. (**D**) Biochemical reconstitution of the recruitment of GFP-TBK1 by mCherry-OPTN to GST-4×Ub coated beads in the presence or absence of increasing amounts of MBP-NAP1. In the indicated wells, unlabeled NDP52 was also added. (top) Diagram of the experimental set-up. (Bottom) Experimental results obtained by confocal imaging. One of three representative experiments is shown. (**E**) Pull-down assay of mCherry-OPTN, mCherry-NDP52, and NAP1 by GFP-TBK1. GFP-TBK1 was pre-loaded onto GFP-Trap beads and then incubated with the protein mixtures as indicated. The relative amounts of mCherry-OPTN, NAP1, and mCherry-NDP52 bound to TBK1 were quantified for the indicated lanes and plotted (right). Data are shown as mean ± s.d. from three independent experiments or as one of three representative Coomassie-stained gels for (E).

We then assessed the extent to which this minimal OPTN peptide could initiate mitophagy, as most of its essential autophagy-driving protein domains had been removed. Yet, treating cells with rapalog for 24 h led to a notable fraction of cells undergoing mitophagy, as demonstrated by the mt-mKeima conversion (**Fig. S6**), albeit to a lesser extent than with full-length OPTN. This observation shows that recruitment of TBK1 is sufficient for mitophagy initiation, consistent with our earlier finding that it can recruit the PI3KC3C1 complex [40].

With this minimal OPTN peptide at hand, we sought to elucidate whether TBK1 recruited through this truncated OPTN axis could enhance NDP52-driven mitophagy. To test this hypothesis, we rescued pentaKO cells with NDP52 and further transduced them with FKBP-OPTN(2-119) and Fis1-FRB. This experimental setup enabled us to measure mitophagy rates by NDP52 upon mitochondrial depolarization by O/A, both in the presence and absence of additional TBK1 recruited through rapalog treatment. While rapalog alone resulted in relatively slow mitophagy activation, displaying only minimal activation from 4 hours onwards, the combined treatment of O/A and rapalog substantially accelerated mitophagy flux (**Fig. 7C** **and S7**). Importantly, this increase in mitophagy flux was not a merely additive effect, based on the kinetics of rapalog treatment alone, especially during the first three hours of treatment where we observed minimal mitophagy induction by rapalog alone, suggesting that the recruitment of TBK1 by OPTN synergistically enhances NDP52-driven mitophagy in cells. This underscores the pivotal role of TBK1 recruitment by OPTN, not only for OPTN’s own function but also for NDP52-mediated mitophagy, as the proximity of TBK1 recruitment by OPTN likely also augments NDP52-mediated mitophagy.

To dissect the interplay between OPTN and NDP52 further, we conducted biochemical reconstitution experiments using agarose beads coated with GST-4×Ub to mimic damaged mitochondrial surfaces. We incubated these beads with OPTN, TBK1, and NAP1 in the presence or absence of NDP52. This confirmed that NAP1 negatively regulates the recruitment of TBK1 towards ubiquitin-bound OPTN in the absence of NDP52, as we showed above (**Fig. 5A**). However, when we added NDP52 to concentrations of NAP1 that would prevent any detectable TBK1 recruitment to the beads, we observed restoration and even a trend towards a slight enhancement of the TBK1 signal on the beads (**Fig. 7D**). To confirm the specificity of the NAP1 sequestration by NDP52, we replaced wild-type NAP1 with a NDP52-binding mutant and observed complete disappearance of TBK1 signal on the ubiquitin-coated beads (**Fig. 7D**). This suggests that the interaction of NAP1 with NDP52 on the cargo allows the mitophagy machinery to overcome the inhibitory effect of NAP1 in terms of TBK1 recruitment during mitophagy initiation.

To assess whether TBK1 indeed converted from binding the OPTN-axis to the NDP52-NAP1/SINTBAD axis, we performed a reverse pull-down experiment using GFP-trap beads coated with EGFP-TBK1. This confirmed that the relative amounts of NAP1 and OPTN determined which of the two cargo receptors, OPTN versus NDP52, was predominately recruited to TBK1 (**Fig. 7E**). These results suggest that NDP52 can support OPTN-driven mitophagy by harnessing NAP1/SINTBAD to recruit further TBK1.

In summary, our findings propose a model in which NAP1/SINTBAD initially set a threshold for mitophagy activation by constraining TBK1 activation via the mitophagy receptor OPTN (**Fig. 8**). This is because OPTN fulfills a primary role in recruiting TBK1 during mitophagy. However, when mitochondrial damage is severe enough, NAP1/SINTBAD transition into a supportive role, acting as cargo co-receptors that bolster NDP52-driven mitophagy. Their sequestration by NDP52 increases TBK1 activation through increased recruitment by OPTN, and this, in return, then boosts NDP52-driven mitophagy again due to the crosstalk from OPTN-TBK1 towards the NDP52-axis, providing an effective feedforward loop once the mitophagy pathway is set in motion.

**Figure 8.**
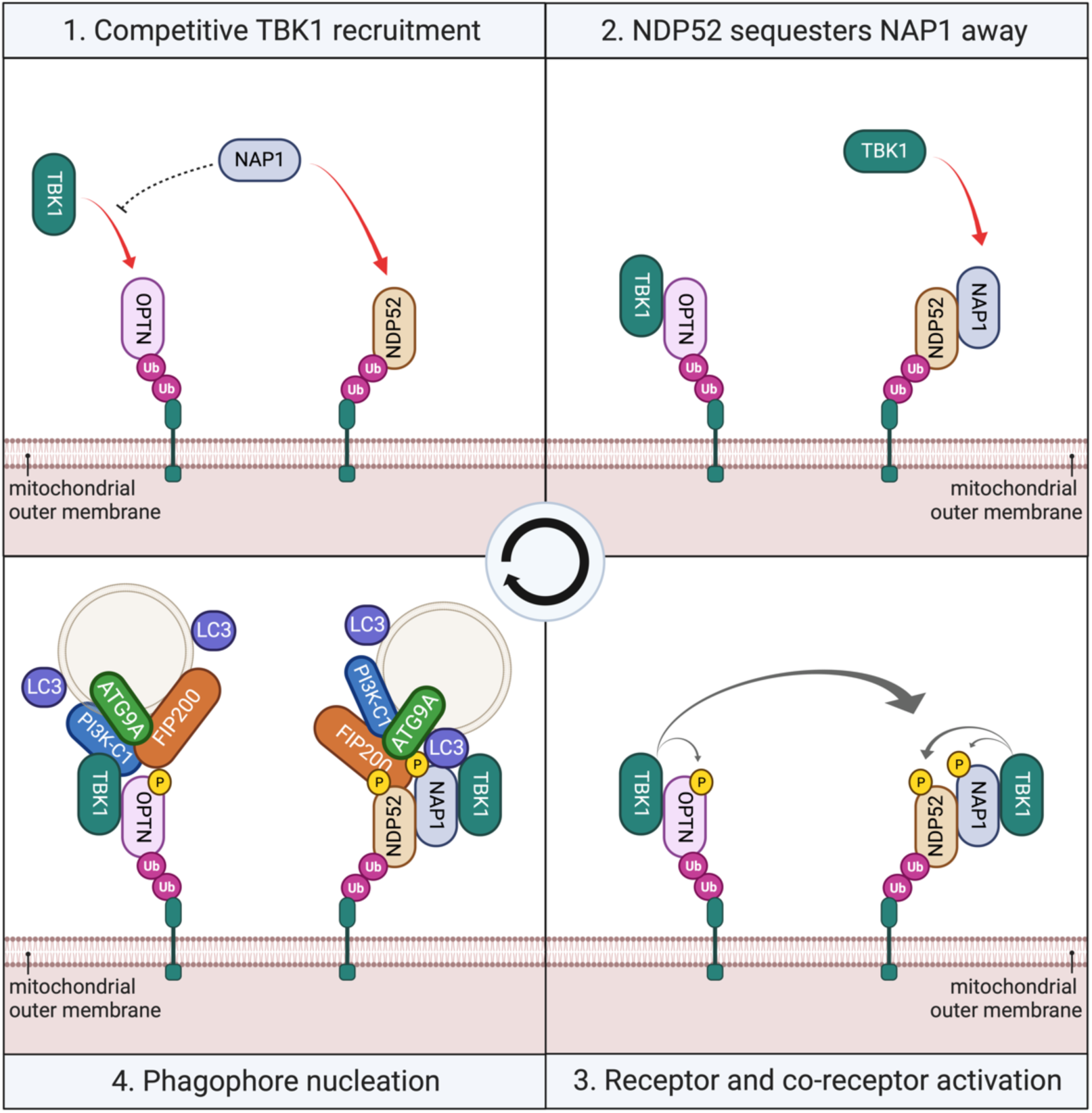
Working model for mitophagy initiation in cells expressing both mitophagy receptors OPTN and NDP52. (**1**) Cargo receptors OPTN and NDP52 are recruited to damaged mitochondria upon accumulation of ubiquitin and phospho-ubiquitin on their surface. OPTN recruits TBK1 but is restricted by NAP1, which competes with OPTN for TBK1-binding. (**2**) However, NDP52 recruits NAP1 to the mitochondrial surface and sequesters NAP1 away, allowing OPTN to recruit and activate more TBK1. (**3**) Clustered TBK1 phosphorylates and activates the cargo receptors and cargo co-receptors, including crosstalk from the OPTN to the NDP52-axis. (**4**) This, in return, facilitates the recruitment of downstream autophagy complexes and the initiation of autophagosome formation.

## DISCUSSION

The regulatory mechanisms that prevent PINK1/Parkin-dependent mitophagy and other selective autophagy pathways from overreacting while ensuring swift progression once initiated are largely elusive. By focusing on the roles of the TBK1 adaptors NAP1/SINTBAD, we uncovered how tightly they are interwoven into this pathway by regulating key activities of the OPTN and NDP52 cargo receptors in completely different ways. In particular, we find that NAP1/SINTBAD act as rheostats, which inhibit mitophagy initiation by restricting recruitment and activation of TBK1 by OPTN, while enhancing NDP52-mediated engulfment of damaged mitochondria.

NAP1/SINTBAD drew our attention due to the central role of TBK1 as a key regulator of selective autophagy pathways and their involvement in supporting NDP52-dependent xenophagy [7, 8, 10]. In addition, we found that NAP1/SINTBAD are recruited and co-degraded with damaged mitochondria (**Fig. 1**). The observation that deletion of NAP1/SINTBAD in wild-type HeLa cells results in acceleration rather than a deceleration of mitophagy (**Fig. 2**) was therefore unexpected. Using cellular and *in vitro* reconstitutions, we dissected how NAP1/SINTBAD interact with NDP52 and OPTN, the key cargo receptors in PINK1/Parkin-dependent mitophagy [30]. In NDP52-driven mitophagy, they exert a stimulatory role (**Fig. 3A-B**), similar to their function in xenophagy, by bridging NDP52 with TBK1 to activate this kinase on the mitochondrial surface. Furthermore, they stabilize the interaction with the core autophagy factor FIP200 (**Fig. 3D**). In contrast, in OPTN-driven mitophagy, NAP1/SINTBAD counteract TBK1 recruitment and activation by directly competing for the same TBK1 binding site (**Fig. 5**). Overall, the inhibitory role of NAP1/SINTBAD seems to prevail, as evidenced by the increased presence of activated TBK1 on damaged mitochondria in the absence of NAP1/SINTBAD in cells expressing both OPTN and NDP52 (**Fig. 5E**).

Our study highlights the central role of TBK1 in coordinating the different mitophagy mechanisms and uncovers an interplay between OPTN-mediated mitophagy and NDP52-mediated mitophagy. This interplay suggests a finely tuned regulation of mitophagy, which may be particularly important for specific cell types. For example, in the brain where NDP52 expression is low and OPTN is the primary mitophagy receptor, competition for TBK1 activation may prevent excessive initiation of mitochondrial degradation. Given the post-mitotic nature of neurons, an excess in mitochondrial degradation could be as detrimental as insufficient activation. This is exemplified by disease-causing mutations in FBXL4, which results in excessive mitochondrial degradation through the NIX/BNIP3 pathway [65–68], and which lead to a severe mitochondrial encephalopathy [69, 70]. Conversely, in cells expressing both mitophagy receptors OPTN and NDP52, NAP1/SINTBAD initially compete with OPTN for TBK1 binding until mitophagy is adequately activated. Subsequently, NAP1/SINTBAD convert into mitophagy-promoting factors by supporting NDP52.

Our results also highlight OPTN’s dominant role in recruiting and activating TBK1 during mitophagy. This is in line with the recent finding that OPTN forms a platform for TBK1 activation from where it engages with TBK1 in a positive feedback loop and observations made with the ALS-causing TBK1-E696K mutant, which has lost its OPTN-binding capacity [39, 60, 61], but not its binding to NAP1 as we show. Although NDP52 can recruit and activate TBK1 in the absence of other cargo receptors, our findings indicate that when OPTN and NDP52 are co-expressed, mitophagy is accelerated when OPTN can more easily recruit TBK1. This hints at a two-tiered mechanism, with OPTN being the first cargo receptor to drive mitophagy at very early stages, followed by NDP52 in a second phase. While the existence of such a mechanism in mitophagy remains in part speculative at this point, previous work has indicated that OPTN and NDP52 are not kinetically interchangeable, with OPTN being more dominant for mitophagy at early time points [39]. Two-step cargo receptor recruitment has also been observed in xenophagy in which NDP52 is recruited initially to invading pathogens via recognition of exposed Galectin 8 molecules [42, 71], subsequently leading to the recruitment of the E3 ubiquitin ligases, such as LUBAC and LRSAM1, which coat the bacterial surface with poly-ubiquitin chains [72–74]. This, in turn, triggers the recruitment of other cargo receptors like OPTN and SQSTM1/p62 [41, 75]. Future research should address whether a similar two-step recruitment mechanism or other diversification mechanisms between cargo receptors underlie our findings.

Additionally, the recent identification of TNIP1 as another mitophagy inhibitor [76] suggests that the inhibitory effect of NAP1/SINTBAD may constitute a more widespread mechanism. TNIP1 was proposed to compete with autophagy receptors for FIP200 binding, which is distinct from how NAP1/SINTBAD inhibit mitophagy, as our results demonstrate that they instead strengthen the NDP52-FIP200 interaction. Nevertheless, identifying these regulatory steps during the early steps of autophagosome biogenesis could offer new therapeutic opportunities, especially in conditions where damaged mitochondria are insufficiently cleared.

In summary, our study uncovers an unexpected additional layer of regulation governing mitophagy initiation and expands our understanding of the complex interplay among various players involved in maintaining mitochondrial quality control. This additional layer may enable cells to respond better to cellular demands and may offer new opportunities for developing new therapeutic strategies aimed at modulating mitophagy in various pathological conditions associated with mitochondrial dysfunction.

## MATERIAL AND METHODS

### Reagents

The following chemicals were used in this study: Oligomycin (A5588, ApexBio), Antimycin A (A8674, Sigma), Q-VD-OPh (A1901, ApexBio), Rapalog A/C hetero-dimerizer (635057, Takara), Bafilomycin A1 (sc-201550, Santa Cruz Biotech), TBK1 inhibitor GSK8612 (S8872, Selleck Chemicals), TBK1 inhibitor BX795 (ENZ-CHM189-0005, Enzo Life Sciences), ULK1/2 inhibitor (MRT68921, BLDpharm), Vps34-IN1 inhibitor (APE-B6179, ApexBio), and DMSO (D2438, Sigma).

### Plasmid Construction

The sequences of all cDNAs were obtained by amplifying existing plasmids, HAP1 cDNA, or through gene synthesis (Genscript). For insect cell expressions, the sequences were codon optimized and gene synthesized (Genscript). With the exception of the NAP1-6×Ala mutant, which was obtained through gene synthesis (Genscript), all other plasmids were generated by Gibson cloning, For Gibson cloning, inserts and vector backbones were generated by PCR amplification or excised from agarose gels after restriction enzyme digestion at 37°C for two hours. The inserts and plasmid backbones were purified with Promega Wizard SV gel and PCR Cleanup System (Promega). Purified inserts and backbones were mixed in a molar 3:1 ratio, respectively, supplemented by a 2× NEBuilder HiFi DNA assembly enzyme mix (New England Biolabs). Gibson reactions were incubated for one hour at 50°C and then transformed into DH5-alpha competent *E. coli* cells. Transformed Gibson reactions were grown overnight on agar plates containing the appropriate selection marker (ampicillin, kanamycin, or chloramphenicol). Single colonies were picked, grown overnight in liquid cultures, and pelleted for DNA plasmid extraction using the GeneJet Plasmid Miniprep kit (Thermo Fisher). The purified plasmid DNA was submitted for DNA Sanger sequencing (MicroSynth AG). All insert sequences were verified by Sanger sequencing. Positive clones were further analyzed by whole plasmid sequencing (Plasmidsaurus). A detailed protocol is available (https://doi.org/10.17504/protocols.io.8epv5x11ng1b/v1).

### Cell lines

All cell lines were cultured at 37°C in humidified 5% C0_2_ atmosphere. HeLa (RRID:CVCL_0058) and HEK293T (RRID:CVCL_0063) cells were acquired from the American Type Culture Collection (ATCC). HAP1 (RRID:CVCL_Y019) cells were purchased from Horizon Discovery. HeLa and HEK293T cells were grown in Dulbecco Modified Eagle Medium (DMEM, Thermo Fisher) supplemented with 10% (v/v) Fetal Bovine Serum (FBS, Thermo Fisher), 25 mM HEPES (15630080, Thermo Fisher), 1% (v/v) non-essential amino acids (NEAA, 11140050, Thermo Fisher), and 1% (v/v) Penicillin-Streptomycin (15140122, Thermo Fisher). HAP1 cells were cultured in Iscove’s Modified Dulbecco’s Medium (IMDM, Thermo Fisher), supplemented with 10% (v/v) Fetal Bovine Serum (FBS, Thermo Fisher) and 1% (v/v) Penicillin-Streptomycin (15140122, Thermo Fisher). All cell lines were tested regularly for mycoplasma contaminations. A detailed protocol is available (https://doi.org/10.17504/protocols.io.n2bvj3y5blk5/v1).

### Generation of CRISPR/Cas9 knockout cells

All knockout cell lines were generated using CRISPR/Cas9. Candidate single-guide RNAs (sgRNAs) were identified using CHOPCHOP (https://chopchop.cbu.uib.no). The sgRNAs were selected to target all common splicing variants. Using Gibson Cloning, the sgRNAs were ordered as short oligonucleotides (Sigma) and cloned into pSpCas9(BB)-2A-GFP vector (RRID:Addgene_48138). The successful insertion of the sgRNAs was verified by Sanger sequencing. A detailed description of this cloning is available (https://doi.org/10.17504/protocols.io.j8nlkkzo6l5r/v1).

Plasmids containing a sgRNA were transfected into HeLa cells with X-tremeGENE8 (Roche). Single GFP-positive cells were sorted by fluorescence-activated cell sorting (FACS) into 96 well plates. Single-cell colonies were expanded and collected for screening to identify positive clones by immunoblotting. Clones that showed a loss of protein expression for the target of interest were further analyzed by Sanger sequencing of the respective genomic regions. After DNA extraction, the regions of interest surrounding the sgRNA target sequence were amplified by PCR and analyzed by Sanger sequencing. The DNA sequences were compared to sequences from the parental line, and the edits were identified using the Synthego ICE v2 CRISPR Analysis Tool. A detailed protocol is available (https://doi.org/10.17504/protocols.io.8epv59yx5g1b/v1).

In cases where we generated multiple gene knockouts in the same cell line, we sequentially transfected sgRNAs for the respective target genes. For NAP1/SINTBAD double knockout clones #13 and #14 (RRID:CVCL_C9DV), the cells were first transfected with NAP1 sgRNA-targeting plasmids, and positive clones were then transfected with SINTBAD sgRNA-targeting plasmids. For NAP1/SINTBAD double knockout clones #20 and #26 in the pentaKO background (RRID:CVCL_C8QB), the pentaKO line (RRID:CVCL_C2VN), first described in Lazarou et al. [30], was transfected with NAP1 and SINTBAD sgRNA-targeting plasmids. For NAP1/SINTBAD/ULK1/2 4KO in the pentaKO background, ULK1/2 were first knocked out in the pentaKO line (RRID:CVCL_C2VS), and this cell line was then used further to delete NAP1/SINTBAD (RRID:CVCL_C9DW).

### Generation of stable cell lines

Stable cell lines were generated using lentiviral or retroviral expression systems. For retroviral transductions, HEK293T cells (RRID:CVCL_0063) were transfected with VSV-G, Gag-Pol, and pBMN constructs containing our gene-of-interest using Lipofectamine 3000 or Lipofectamine LTX (L3000008 or A12621, Thermo Fisher). The next day, the medium was exchanged with fresh media. Viruses were harvested 48 h and 72 h after transfection. The retrovirus-containing supernatant was collected and filtered to avoid cross-over of HEK293T cells into our HeLa cultures. HeLa cells, seeded at a density of 800k per well, were infected by the retrovirus-containing supernatant in the presence of 8 mg/ml polybrene (Sigma) for 24h. The infected HeLa cells were expanded, and 10 days after infection, they were sorted by FACS to match equal expression levels where possible. A detailed protocol is available (https://doi.org/10.17504/protocols.io.81wgbyez1vpk/v1).

The following retroviral vectors were used in this study: pBMN-HA-NAP1 (RRID:Addgene_208868), pBMN-HA-NAP1 delta-TBK1 (L226Q/L233Q) (RRID:Addgene_208869), pBMN-HA-SINTBAD (RRID:Addgene_), pBMN-mEGFP-OPTN (RRID:Addgene_188784), pBMN-mEGFP-NDP52 (RRID:Addgene_188785), pBMN-BFP-Parkin (RRID:Addgene_186221), and pCHAC-mito-mKeima (RRID:Addgene_72342). Empty backbones used to generate these retroviral vectors were pBMN-HA-C1 (RRID:Addgene_188645), pBMN-mEGFP (RRID:Addgene_188643), and pBMN-BFP-C1 (RRID:Addgene_188644).

For lentiviral transductions, HEK293T cells (RRID:CVCL_0063) were transfected with VSV-G, Gag-Pol, and pHAGE constructs containing our gene-of-interest using Lipofectamine 3000 (L3000008, Thermo Fisher). The next day, the medium was exchanged with fresh media. Viruses were harvested 48 h and 72 h after transfection. The lentivirus-containing supernatant was collected and filtered to avoid cross-over of HEK293T cells into our HeLa cultures. HeLa cells, seeded at a density of 800k per well, were infected by the lentivirus-containing supernatant in the presence of 8 mg/ml polybrene (Sigma) for 24 h. The infected HeLa cells were expanded, and 10 days after infection, they were used for experiments. A detailed protocol is available (https://doi.org/10.17504/protocols.io.6qpvr3e5pvmk/v1).

The following lentiviral vectors were used in this study: pHAGE-FKBP-GFP-NDP52 (RRID:Addgene_135296), pHAGE-FKBP-GFP-NAP1 (RRID:Addgene_208862), pHAGE-FKBP-GFP-NAP1 delta-NDP52 (S37K/A44E) (RRID:Addgene_208863), pHAGE-FKBP-GFP-NAP1 delta-FIP200 (I11S/L12S) (RRID:Addgene_208864), pHAGE-FKBP-GFP-NAP1 delta-TBK1 (L226Q/L233Q) (RRID:Addgene_208865), pHAGE-FKBP-GFP-OPTN (RRID:Addgene_208866), pHAGE-FKBP-GFP-OPTN (2-119) (RRID:Addgene_208867), pHAGE-mt-mKeima-P2A-FRB-Fis1 (RRID:Addgene_135295).

### Mitophagy experiments

To induce mitophagy, cells were treated with 10 µM Oligomycin (A5588, ApexBio) and 4 µM Antimycin A (A8674, Sigma). In case cells were treated for more than 8 h, we also added 10 µM Q-VD-OPh (A1901, ApexBio) to suppress apoptosis. Samples were then analyzed by SDS-PAGE and western blot or flow cytometry. A detailed protocol is available (https://doi.org/10.17504/protocols.io.n2bvj3yjnlk5/v1).

### Nutrient starvation experiments

To induce bulk autophagy, cells were starved by culturing them in Hank balanced salt medium (HBSS, Thermo Fisher). Cells were collected and analyzed by SDS-PAGE and western blot analysis. A detailed protocol is available (https://doi.org/10.17504/protocols.io.4r3l228b3l1y/v1).

### Rapalog-induced chemical dimerization experiments

The chemical-induced dimerization (CID) experiments were performed using the FRB-Fis1 and FKBP fused to our gene of interest system. After consecutive lentiviral transduction of HeLa cells with both constructs, in which the FRB-Fis1 also expresses mitochondrially targeted monoKeima (mt-mKeima), cells were treated with the Rapalog A/C hetero-dimerizer rapalog (635057, Takara) for 24 h. Cells were then analyzed by flow cytometry. A detailed protocol is available (https://doi.org/10.17504/protocols.io.n92ldmyynl5b/v1).

### Flow cytometry

For mitochondrial flux experiments, 800K cells were seeded in 6 well plates one day before the experiment. Mitophagy was induced by treating the cells for the indicated times with a cocktail of oligomycin and antimycin A (O/A), as described above. Cells were collected by removing the medium, washing the cells with 1× PBS (14190169, Thermo Fisher), trypsinization (T3924, Sigma), and resuspending in complete DMEM medium (41966052, Thermo Fisher). Filtered through 35 µm cell-strainer caps (352235, Falcon) and analyzed by an LSR Fortessa Cell Analyzer (BD Biosciences). Lysosomal mt-mKeima was measured using dual excitation ratiometric pH measurements at 405 (pH 7) and 561 (pH 4) nm lasers with 710/50-nm and 610/20-nm detection filters, respectively. Additional channels used for fluorescence compensation were BFP and GFP. Single fluorescence vector expressing cells were prepared to adjust photomultiplier tube voltages to make sure the signal was within detection limits, and to calculate the compensation matrix in BD FACSDiva Software. Depending on the experiment, we gated for BFP-positive, GFP-positive, and mKeima-positive cells with the appropriate compensation. For each sample, 10,000 mKeima-positive events were collected, and data were analyzed in FlowJo (version 10.9.0). Our protocol was based on the previously described protocol (https://doi.org/10.17504/protocols.io.q26g74e1qgwz/v1).

For Rapalog-induced mitophagy experiments, cells were seeded as described above and treated for 24 h with 500 nM Rapalog A/C hetero-dimerizer (Takara). Cells were collected as described above, and the mt-mKeima ratio (561/405) was quantified by an LSR Fortessa Cell Analyzer (BD Biosciences). The cells were gated for GFP/mt-mKeima double-positive cells. Data were analyzed using FlowJo (version 10.9.0). A detailed protocol is available (https://doi.org/10.17504/protocols.io.n92ldmyynl5b/v1).

### Cellular fractionation and mitochondrial isolation

HeLa cells were seeded in 15 cm dishes and grown until confluence. Cells were treated with DMSO or O/A for the indicated time. Mitochondria were isolated as described previously [77]. In brief, cells were collected by trypsinization, centrifuged at 300*g* for 5 min at 4°C, and the cell pellet was washed in PBS to remove the remaining medium. A fraction of the PBS-washed cell pellet was transferred to a new tube and lysed in RIPA buffer (50 mM Tris-HCl pH 8.0, 150 mM NaCl, 0.5% sodium deoxycholate, 0.1% SDS, 1% NP-40) supplemented by cOmplete EDTA-free protease inhibitors (11836170001, Roche) and phosphatase inhibitors (Phospho-STOP, 4906837001, Roche). This sample served as a whole cell lysate (WCL) reference. The remaining PBS-washed cells were processed further for mitochondrial isolation. In this case, the PBS was removed from the cell pellet, and the cells were resuspended in 1 ml mitochondrial isolation buffer (250 mM mannitol, 0.5 mM EGTA, and 5 mM HEPES-KOH pH 7.4). The cells were lysed by 15 strokes with 26.5 G needle (303800, Becton Dickinson). The homogenate was then centrifuged twice at 600*g* for 10 min at 4°C to pellet cell debris, nuclei, and intact cells. The supernatant was collected and centrifuged twice at 7,000*g* for 10 min at 4°C to pellet mitochondria. The supernatant was removed, and the mitochondrial pellet was resuspended in 1 ml of mitochondrial isolation buffer. The resuspended mitochondrial pellets were centrifuged two more times at 10,000*g* for 10 min at 4°C. After removal of the supernatant, the pellets were resuspended in the mitochondrial isolation buffer. The final mitochondrial pellet was lysed in RIPA buffer and processed further for western blot analysis. A detailed protocol is available (https://doi.org/10.17504/protocols.io.kqdg3x4zzg25/v1).

### SDS-PAGE and western blot analysis

For SDS-PAGE and western blot analysis, we collected cells by trypsinization and subsequent centrifugation at 300*g* for 5 min at 4°C. Cell pellets were washed in PBS and centrifuged once more at 300*g* for 5 min at 4°C. The supernatant was removed and the cell pellets were lysed in RIPA buffer (50 mM Tris-HCl pH 8.0, 150 mM NaCl, 0.5% sodium deoxycholate, 0.1% SDS, 1% NP-40) supplemented by cOmplete EDTA-free protease inhibitors (11836170001, Roche) and phosphatase inhibitors (Phospho-STOP, 4906837001, Roche). After incubating in RIPA buffer for 20 min on ice, samples were cleared by centrifugation at 20,000*g* for 10 min at 4°C. The soluble supernatant fraction was collected and protein concentrations were measured using the Pierce Detergent Compatible Bradford Assay Kit (23246, Thermo Fisher). Samples were then adjusted for equal loading and mixed with 6× protein loading dye, supplemented with 100 mM DTT, and boiled for 5 min at 95°C. Samples were loaded on 4-12% SDS-PAGE gels (NP0321BOX, NP0322BOX, or NP0323BOX, Thermo Fisher) with PageRuler Prestained protein marker (Thermo Fisher). Proteins were transferred onto nitrocellulose membranes (RPN132D, GE Healthcare) for 1 h at 4°C using the Mini Trans-Blot Cell (Bio-Rad). After the transfer, membranes were blocked with 5% milk powder dissolved in PBS-Tween (0.1% Tween 20) for 1 h at room temperature. The membranes were incubated overnight at 4°C with primary antibodies dissolved in the blocking buffer, washed three times for 5 min, and incubated with species-matched secondary horseradish peroxidase (HRP)-coupled antibodies diluted 1:10,000 in blocking buffer for 1 h at room temperature. Membranes were afterward washed three times with PBS-T and processed further for western blot detection. Membranes were incubated with SuperSignal West Femto Maximum Sensitivity Substrate (34096, Thermo Fisher) and imaged with a ChemiDoc MP Imaging system (Bio-Rad). Images were analyzed with ImageJ [78]. A detailed protocol is available (https://doi.org/10.17504/protocols.io.eq2lyj33plx9/v1).

The primary antibodies used in this study are: anti-4E-BP1 (1:1000, Proteintech Cat# 60246-1-Ig, RRID:AB_2881368), anti-ATG5 (1:1000, Cell Signaling Technology Cat# 12994, RRID:AB_2630393), anti-β-Actin (1:5000, Abcam Cat# ab20272, RRID:AB_445482), anti-COXII (1:1000, Abcam Cat# ab110258, RRID:AB_10887758) or (1:1000, Cell Signaling Technology Cat# 31219, RRID:AB_2936222), anti-FIP200 (1:1000, Cell Signaling Technology Cat# 12436, RRID:AB_2797913), anti-HA (1:500, Cell Signaling Technology Cat# 2367, RRID:AB_10691311) or (1:1000, Cell Signaling Technology Cat# 3724, RRID:AB_1549585), anti-mHSP60 (1:1000, Abcam Cat# ab128567, RRID:AB_11145464), anti-NAP1 (1:1000, Abcam Cat# ab192253, RRID:AB_2941051), anti-p62/SQSTM1 (1:1000, Abnova Cat# H00008878-M01, RRID:AB_437085), anti-SINTBAD (1:1000, Cell Signaling Technology Cat# 8605, RRID:AB_10839270), anti-TBK1 (1:1000, Cell Signaling Technology Cat# 38066, RRID:AB_2827657) or (1:1000, Cell Signaling Technology Cat# 3013, RRID:AB_2199749), anti-phospho-TBK1 S172 (1:1000, Cell Signaling Technology Cat# 5483, RRID:AB_10693472), anti-α-Tubulin (1:5000, Abcam Cat# ab7291, RRID:AB_2241126), anti-phospho-Ubiquitin S65 (1:2000, Millipore Cat# ABS1513-I, RRID:AB_2858191), anti-ULK1 (1:1000, Cell Signaling Technology Cat# 8054, RRID:AB_11178668).

The secondary antibodies used in this study are: HRP conjugated polyclonal goat anti-mouse (Jackson ImmunoResearch Labs Cat# 115-035-003, RRID:AB_10015289), HRP conjugated polyclonal goat anti-rabbit (Jackson ImmunoResearch Labs Cat# 111-035-003, RRID:AB_2313567).

### Immunofluorescence and confocal microscopy

Cells were seeded on HistoGrip (Thermo Fisher) coated glass coverslips in 24 well plates. Cells were allowed to adhere overnight and treated as indicated prior to fixation. Cells were fixed in 4% (w/v) paraformaldehyde (PFA), diluted in 100 mM phosphate buffer, for 10 min at room temperature. The PFA was removed, and samples were washed three times with 1× PBS. Cells were permeabilized with 0.1% (v/v) Triton X-100, diluted in 1× PBS, for 10 min. After permeabilization, samples were blocked for 15 min with 3% (v/v) goat serum diluted in 1× PBS with 0.1% Triton X-100. Primary antibodies, diluted in blocking buffer, were incubated with the samples for 90 min. Unbound antibodies were removed in three washing steps with 1× PBS. Secondary antibodies, diluted in blocking buffer, were incubated with the samples for 60 min. Secondary antibodies were conjugated to AlexaFluor-488, Alexa-Fluor-555, Alexa-Fluor-633, or Alexa-Fluor-647 (Thermo Fisher). Unbound secondary antibodies were removed by three washes with 1× PBS before coverslips were mounted onto glass slides with DABCO-glycerol mounting medium. Coverslips were imaged with an inverted Leica SP8 confocal laser scanning microscope equipped with an HC Plan Apochromat CS2 63×/1.40 oil immersion objective (Leica Microsystems). Images were acquired in three dimensions using z-stacks, with a minimum range of 1.8 µM and a maximum voxel size of 90 nm laterally (x,y) and 300 nm axially (z), using a Leica HyD Hybrid detector (Leica Microsystems) and the Leica Application Suite X (LASX v2.0.1). The z-stack images are displayed as maximum-intensity projections. Three images were taken for each sample. A detailed protocol is available (https://doi.org/10.17504/protocols.io.5qpvobz99l4o/v1).

### Protein expression and purification

Linear tetra-ubiquitin fused to GST (GST-4×Ub) was cloned into a pGEX-4T1 vector (RRID:Addgene_199779). After the transformation of the pGEX-4T1 vector encoding GST-4×Ub in *E. coli* Rosetta pLySS cells, cells were grown in LB medium at 37°C until an OD_600_ of 0.4 and then continued at 18°C. Once the cells reached an OD_600_ of 0.8, protein expression was induced with 100 µM isopropyl β-D-1-thiogalactopyranoside (IPTG) for 16 h at 18°C. Cells were collected by centrifugation and resuspended in lysis buffer (50 mM Tris-HCl pH 7.4, 300 mM NaCl, 2 mM MgCl_2_, 1 mM DTT, cOmplete EDTA-free protease inhibitors (Roche), and DNase (Sigma)). Cell lysates were sonicated twice for 30 s. Lysates were cleared by centrifugation at 18,000 rpm for 45 min at 4°C in a SORVAL RC6+ centrifuge with an F21S-8×50Y rotor (Thermo Scientific). The supernatant was collected and incubated with pre-equilibrated Glutathione Sepharose 4B beads (GE Healthcare) for 2 h at 4°C with gentle shaking to bind GST-4×Ub. Samples were centrifuged to pellet the beads and remove the unbound lysate. Beads were then washed twice with wash buffer (50 mM Tris-HCl pH 7.4, 300 mM NaCl, 1 mM DTT), once with high salt wash buffer (50 mM Tris-HCl pH 7.4, 700 mM NaCl, 1 mM DTT), and two more times with wash buffer (50 mM Tris-HCl pH 7.4, 300 mM NaCl, 1 mM DTT). Beads were incubated overnight with 4 ml of 50 mM reduced glutathione dissolved in wash buffer (50 mM Tris-HCl pH 7.4, 300 mM NaCl, 1 mM DTT) at 4°C, to elute GST-4×Ub from the beads. To collect the supernatant, the beads were collected by centrifugation. The beads were washed twice with 4 ml of wash buffer, and the supernatant was collected. The supernatant fractions were pooled, filtered through a 0.45 µm syringe filter, concentrated with 10 kDa cut-off Amicon filter (Merck Millipore), and loaded onto a pre-equilibrated Superdex 200 Increase 10/300 GL column (Cytiva). Proteins were eluted with SEC buffer (25 mM Tris-HCl pH 7.4, 150 mM NaCl, 1 mM DTT). Fractions were analyzed by SDS-PAGE and Coomassie staining. Fractions containing purified GST-4×Ub were pooled. After concentrating the purified protein, the protein was aliquoted and snap-frozen in liquid nitrogen. Proteins were stored at –80°C. A detailed protocol is available (https://doi.org/10.17504/protocols.io.q26g7pbo1gwz/v1).

For mCherry-OPTN, we cloned human OPTN cDNA in a pETDuet-1 vector with an N-terminal 6×His tag followed by a TEV cleavage site (RRID:Addgene_190191). After the transformation of the pETDuet-1 vector encoding 6×His-TEV-mCherry-OPTN in *E. coli* Rosetta pLySS cells, cells were grown in 2×TY medium at 37°C until an OD_600_ of 0.4 and then continued at 18°C. Once the cells reached an OD_600_ of 0.8, protein expression was induced with 50 µM IPTG for 16 h at 18°C. Cells were collected by centrifugation and resuspended in lysis buffer (50 mM Tris-HCl pH 7.4, 300 mM NaCl, 2 mM MgCl_2_, 5% glycerol, 10 mM Imidazole, 2 mM β-mercaptoethanol, cOmplete EDTA-free protease inhibitors (Roche), CIP protease inhibitor (Sigma), and DNase (Sigma)). Cell lysates were sonicated twice for 30 s. Lysates were cleared by centrifugation at 18,000 rpm for 45 min at 4°C in a SORVAL RC6+ centrifuge with an F21S-8×50Y rotor (Thermo Scientific). The supernatant was filtered through an 0.45 µm filter and loaded onto a pre-equilibrated 5 ml His-Trap HP column (Cytiva). After His tagged proteins were bound to the column, the column was washed with three column volumes of wash buffer (50 mM Tris-HCl pH 7.4, 300 mM NaCl, 10 mM Imidazole, 2 mM β-mercaptoethanol). Proteins were then eluted with a stepwise imidazole gradient (30, 75, 100, 150, 225, 300 mM). Fractions at 75-100 mM imidazole contained the 6×His-TEV-mCherry-OPTN and were pooled. The pooled samples were incubated overnight with TEV protease at 4°C. After the 6×His tag was cleaved off, the protein was concentrated using a 50 kDa cut-off Amicon filter (Merck Millipore) and loaded onto a pre-equilibrated Superdex 200 Increase 10/300 GL column (Cytiva). Proteins were eluted with SEC buffer (25 mM Tris-HCl pH 7.4, 150 mM NaCl, 1 mM DTT). Fractions were analyzed by SDS-PAGE and Coomassie staining.

Fractions containing purified mCherry-OPTN were pooled. After concentrating the purified protein, the protein was aliquoted and snap-frozen in liquid nitrogen. Proteins were stored at –80°C. A detailed protocol is available (https://doi.org/10.17504/protocols.io.4r3l225djl1y/v1). For mCherry-NDP52, we cloned human NDP52 cDNA in a pETDuet-1 vector with an N-terminal 6×His tag followed by a TEV cleavage site (RRID:Addgene_187829). After the transformation of the pETDuet-1 vector encoding 6×His-TEV-mCherry-NDP52 in *E. coli* Rosetta pLySS cells, cells were grown in 2×TY medium at 37°C until an OD_600_ of 0.4 and then continued at 18°C. Once the cells reached an OD_600_ of 0.8, protein expression was induced with 50 µM IPTG for 16 h at 18°C. Cells were collected by centrifugation and resuspended in lysis buffer (50 mM Tris-HCl pH 7.4, 300 mM NaCl, 2 mM MgCl_2_, 5% glycerol, 10 mM Imidazole, 2 mM β-mercaptoethanol, cOmplete EDTA-free protease inhibitors (Roche), CIP protease inhibitor (Sigma), and DNase (Sigma)). Cell lysates were sonicated twice for 30 s. Lysates were cleared by centrifugation at 18,000 rpm for 45 min at 4°C in a SORVAL RC6+ centrifuge with an F21S-8×50Y rotor (Thermo Scientific). The supernatant was filtered through an 0.45 µm filter and loaded onto a pre-equilibrated 5 ml His-Trap HP column (Cytiva). After His tagged proteins were bound to the column, the column was washed with three column volumes of wash buffer (50 mM Tris-HCl pH 7.4, 300 mM NaCl, 10 mM Imidazole, 2 mM β-mercaptoethanol). Proteins were then eluted with a stepwise imidazole gradient (30, 75, 100, 150, 225, 300 mM). Fractions at 75-100 mM imidazole contained the 6×His-TEV-mCherry-NDP52 and were pooled. The pooled samples were incubated overnight with TEV protease at 4°C. After the 6×His tag was cleaved off, the protein was concentrated using a 50 kDa cut-off Amicon filter (Merck Millipore) and loaded onto a pre-equilibrated Superdex 200 Increase 10/300 GL column (Cytiva). Proteins were eluted with SEC buffer (25 mM Tris-HCl pH 7.4, 150 mM NaCl, 1 mM DTT). Fractions were analyzed by SDS-PAGE and Coomassie staining. Fractions containing purified mCherry-NDP52 were pooled. After concentrating the purified protein, the protein was aliquoted and snap-frozen in liquid nitrogen. Proteins were stored at –80°C. A detailed protocol is available (https://doi.org/10.17504/protocols.io.5qpvobdr9l4o/v1). Human NDP52 cDNA was cloned into a pGST2 vector with an N-terminal GST tag followed by a TEV cleavage site (RRID:Addgene_187828). After the transformation of the pGST2 vector encoding GST-TEV-NDP52 in *E. coli* Rosetta pLySS cells, cells were grown in 2×TY medium at 37°C until an OD_600_ of 0.4 and then continued at 18°C. Once the cells reached an OD_600_ of 0.8, protein expression was induced with 50 µM IPTG for 16 h at 18°C. Cells were collected by centrifugation and resuspended in lysis buffer (50 mM Tris-HCl pH 7.4, 300 mM NaCl, 1 mM DTT, 2 mM MgCl_2_, 2 mM β-mercaptoethanol, cOmplete EDTA-free protease inhibitors (Roche), and DNase (Sigma)). Cell lysates were sonicated twice for 30 s. Lysates were cleared by centrifugation at 18,000 rpm for 45 min at 4°C in a SORVAL RC6+ centrifuge with an F21S-8×50Y rotor (Thermo Scientific). The supernatant was collected and incubated with pre-equilibrated Glutathione Sepharose 4B beads (GE Healthcare) for 2 h at 4°C with gentle shaking to bind GST-NDP52. Samples were centrifuged to pellet the beads and remove the unbound lysate. Beads were then washed twice with wash buffer (50 mM Tris-HCl pH 7.4, 300 mM NaCl, 1 mM DTT), once with high salt wash buffer (50 mM Tris-HCl pH 7.4, 700 mM NaCl, 1 mM DTT), and two more times with wash buffer (50 mM Tris-HCl pH 7.4, 300 mM NaCl, 1 mM DTT). Beads were incubated overnight with TEV protease at 4°C. After the GST tag was cleaved off, the protein was filtered through a 0.45 µm syringe filter, concentrated using a 30 kDa cut-off Amicon filter (Merck Millipore), and loaded onto a pre-equilibrated Superdex 200 Increase 10/300 GL column (Cytiva). Proteins were eluted with SEC buffer (25 mM Tris-HCl pH 7.4, 150 mM NaCl, 1 mM DTT). Fractions were analyzed by SDS-PAGE and Coomassie staining. Fractions containing purified NDP52 were pooled. After concentrating the purified protein, the protein was aliquoted and snap-frozen in liquid nitrogen. Proteins were stored at –80°C. A detailed protocol is available (https://doi.org/10.17504/protocols.io.36wgq35xklk5/v1).

To purify NAP1 or GST-NAP1, human NAP1 cDNA was synthesized and cloned in a pcDNA3.1 vector (Genscript), from where it was subcloned into a pGEX-4T1 vector with an N-terminal GST tag followed by a TEV cleavage site (RRID:Addgene_208870). For expression of unlabeled NAP1 in *E. coli* (which we used in Figure 5B, Figure 7E, and Figure S5) or GST-NAP1 (which we used in Figure 6E and Figure S3) the pGEX-4T1 vector encoding GST-TEV-NAP1 was transformed into *E. coli* Rosetta pLySS cells, cells were grown in 2×TY medium at 37°C until an OD_600_ of 0.4 and then continued at 18°C. Once the cells reached an OD_600_ of 0.8, protein expression was induced with 50 µM IPTG for 16 h at 18°C. Cells were collected by centrifugation and resuspended in lysis buffer (50 mM Tris-HCl pH 7.4, 300 mM NaCl, 1 mM DTT, 2 mM MgCl_2_, 5% glycerol, 2 mM β-mercaptoethanol, cOmplete EDTA-free protease inhibitors (Roche), and DNase (Sigma)). Cell lysates were sonicated twice for 30 s. Lysates were cleared by centrifugation at 18,000 rpm for 45 min at 4°C in a SORVAL RC6+ centrifuge with an F21S-8×50Y rotor (Thermo Scientific). The supernatant was collected and incubated with pre-equilibrated Glutathione Sepharose 4B beads (GE Healthcare) for 2 h at 4°C with gentle shaking to bind GST-TEV-NAP1. Samples were centrifuged to pellet the beads and remove the unbound lysate. Beads were then washed twice with wash buffer (50 mM Tris-HCl pH 7.4, 300 mM NaCl, 5% glycerol, 1 mM DTT), once with high salt wash buffer (50 mM Tris-HCl pH 7.4, 700 mM NaCl, 5% glycerol, 1 mM DTT), and two more times with wash buffer (50 mM Tris-HCl pH 7.4, 300 mM NaCl, 5% glycerol, 1 mM DTT). Beads were incubated overnight at 4°C with TEV protease or 4 ml of 50 mM reduced glutathione dissolved in wash buffer (50 mM Tris-HCl pH 7.4, 300 mM NaCl, 5% glycerol, 1 mM DTT). After the proteins were released from the beads, the GST-NAP1 protein was filtered through a 0.45 µm syringe filter, concentrated using a 30 kDa cut-off Amicon filter (Merck Millipore), or 10 kDa cut-off in case of unlabeled NAP1, and loaded onto a pre-equilibrated Superose 6 Increase 10/300 GL column (Cytiva). Proteins were eluted with SEC buffer (25 mM Tris-HCl pH 7.4, 300 mM NaCl, 1 mM DTT). Fractions were analyzed by SDS-PAGE and Coomassie staining. Fractions containing purified NAP1 or GST-NAP1 protein were pooled. After concentrating the purified protein, the protein was aliquoted and snap-frozen in liquid nitrogen. Proteins were stored at –80°C. A detailed protocol can be found here (https://doi.org/10.17504/protocols.io.kqdg3xk41g25/v1).

To purify MBP-NAP1, human NAP1 cDNA was gene-synthesized (by Genscript) and subcloned into a pGEX-4T1 vector with an N-terminal MBP-tag followed by a TEV cleavage site before wild-type NAP1 (RRID:Addgene_208871), NAP1 delta-NDP52 (S37K/A44E) (RRID:Addgene_208872), or NAP1 delta-TBK1 (L226Q/L233Q) (RRID:Addgene_208873). For expression of MBP-TEV-NAP1 in *E. coli* (which we used for Figure 7D), the pGEX-4T1 vector encoding MBP-TEV-NAP1 was transformed into *E. coli* Rosetta pLySS cells, cells were grown in 2xTY medium at 37°C until an OD_600_ of 0.4 and then continued at 18°C. Once the cells reached an OD_600_ of 0.8, protein expression was induced with 50 µM IPTG for 16 h at 18°C. Cells were collected by centrifugation and resuspended in lysis buffer (50 mM Tris-HCl pH 7.4, 300 mM NaCl, 1 mM DTT, 2 mM MgCl_2_, 5% glycerol, 2 mM β-mercaptoethanol, cOmplete EDTA-free protease inhibitors (Roche), and DNase (Sigma)). Cell lysates were sonicated twice for 30 s and then cleared by centrifugation at 18,000 rpm for 45 min at 4°C in a SORVAL RC6+ centrifuge with an F21S-8×50Y rotor (Thermo Scientific). The supernatant was collected and incubated with pre-equilibrated Amylose beads (Biolabs) for 2 h at 4°C with gentle shaking to bind MBP-TEV-NAP1. Samples were centrifuged to pellet the beads and remove the unbound lysate. Beads were then washed twice with wash buffer (50 mM Tris-HCl pH 7.4, 300 mM NaCl, 5% glycerol, 1 mM DTT), once with high salt wash buffer (50 mM Tris-HCl pH 7.4, 700 mM NaCl, 5% glycerol, 1 mM DTT), and two more times with wash buffer (50 mM Tris-HCl pH 7.4, 300 mM NaCl, 5% glycerol, 1 mM DTT). Beads were incubated overnight at 4°C with 250 mM D-maltose (Santa Cruz) dissolved in wash buffer (50 mM Tris-HCl pH 7.4, 300 mM NaCl, 5% glycerol, 1 mM DTT). After the proteins were released from the beads, the MBP-TEV-NAP1 protein was filtered through a 0.45 µm syringe filter, concentrated using a 30 kDa cut-off Amicon filter (Merck Millipore), and loaded onto a pre-equilibrated Superose 6 Increase 10/300 GL column (Cytiva). Proteins were eluted with SEC buffer (25 mM Tris-HCl pH 7.4, 300 mM NaCl, 1 mM DTT). Fractions were analyzed by SDS-PAGE and Coomassie staining. Fractions containing purified MBP-TEV-NAP1 protein were pooled. After concentrating the purified protein, the protein was aliquoted and snap-frozen in liquid nitrogen. Proteins were stored at –80°C. A detailed protocol can be found here (https://doi.org/10.17504/protocols.io.ewov1q2ykgr2/v1).

To purify SINTBAD-GFP and SINTBAD-mCherry from insect cells, we purchased gene-synthesized codon-optimized GST-TEV-SINTBAD-EGFP and GST-TEV-SINTBAD-mCherry in a pFastBac-Dual vector from Genscript (RRID:Addgene_198035 and RRID:Addgene_208874). The constructs were used to generate bacmid DNA, using the Bac-to-Bac system, by amplification in DH10BacY cells [79]. After the bacmid DNA was verified by PCR for insertion of the transgene, we purified bacmid DNA for transfection into Sf9 insect cells (12659017, Thermo Fisher, RRID:CVCL_0549). To this end, we mixed 2500 ng of plasmid DNA with FuGene transfection reagent (Promega) and transfected 1 million Sf9 cells seeded in a 6 well plate. About 7 days after transfection, the V0 virus was harvested and used to infect 40 ml of 1 million cells per ml of Sf9 cells. The viability of the cultures was closely monitored and upon the decrease in viability and confirmation of yellow fluorescence, we collected the supernatant after centrifugation and stored this as V1 virus. For expressions, we infected 1 L of Sf9 cells, at 1 million cells per ml, with 1 ml of V1 virus. When the viability of the cells decreased to 90-95%, cells were collected by centrifugation. Cell pellets were washed with 1× PBS and flash-frozen in liquid nitrogen. Pellets were stored at –80°C. For purification of SINTBAD-GFP and SINTBAD-mCherry, pellets were resuspended in 25 ml lysis buffer (50 mM Tris-HCl pH 7.4, 300 mM NaCl, 1 mM DTT, 2 mM MgCl_2_, 5% glycerol, 2 mM β-mercaptoethanol, 1 µl benzonase (Sigma), cOmplete EDTA-free protease inhibitors (Roche), CIP protease inhibitor (Sigma)). Cells were homogenized with a douncer. Cell lysates were cleared by centrifugation at 18,000 rpm for 45 min at 4°C in a SORVAL RC6+ centrifuge with an F21S-8×50Y rotor (Thermo Scientific). The supernatant was collected and incubated with pre-equilibrated Glutathione Sepharose 4B beads (GE Healthcare) for 2 h at 4°C with gentle shaking to bind GST-TEV-SINTBAD-EGFP or GST-TEV-SINTBAD-mCherry. Samples were centrifuged to pellet the beads and remove the unbound lysate. Beads were then washed twice with wash buffer (50 mM Tris-HCl pH 7.4, 300 mM NaCl, 5% glycerol, 1 mM DTT), once with high salt wash buffer (50 mM Tris-HCl pH 7.4, 700 mM NaCl, 5% glycerol, 1 mM DTT), and two more times with wash buffer (50 mM Tris-HCl pH 7.4, 300 mM NaCl, 5% glycerol, 1 mM DTT). Beads were incubated overnight with TEV protease in wash buffer (50 mM Tris-HCl pH 7.4, 300 mM NaCl, 5% glycerol, 1 mM DTT) at 4°C. After the proteins were released from the beads by the TEV protease, the supernatant was collected after centrifugation of the beads. The beads were washed twice with 4 ml of wash buffer, and the supernatant was collected. The supernatant fractions were pooled, filtered through a 0.45 µm syringe filter, and concentrated with a 30 kDa cut-off Amicon filter (Merck Millipore). The proteins were loaded onto a pre-equilibrated Superdex 200 Increase 10/300 GL column (Cytiva). Proteins were eluted with SEC buffer (25 mM Tris-HCl pH 7.4, 300 mM NaCl, 1 mM DTT). Fractions were analyzed by SDS-PAGE and Coomassie staining. Fractions containing purified SINTBAD-GFP and SINTBAD-mCherry were pooled. After concentrating the purified protein, the protein was aliquoted and snap-frozen in liquid nitrogen. Proteins were stored at –80°C. A detailed protocol can be found here (https://doi.org/10.17504/protocols.io.rm7vzb1o8vx1/v1).

To purify TBK1 and GFP-TBK1, we purchased gene-synthesized codon-optimized GST-TEV-TBK1 and GST-TEV-EGFP-TBK1 in a pFastBac-Dual vector from Genscript (RRID:Addgene_208875 and Addgene_187830) for expression in insect cells. The V1 virus was generated as described above for SINTBAD. For expressions, we infected 1 L of Sf9 cells (12659017, Thermo Fisher, RRID:CVCL_0549), at 1 million cells per ml, with 1 ml of V1 virus. When the viability of the cells decreased to 90-95%, cells were collected by centrifugation. Cell pellets were washed with 1× PBS and flash-frozen in liquid nitrogen. Pellets were stored at –80°C. For purification of SINTBAD-GFP and SINTBAD-mCherry, pellets were resuspended in 25 ml lysis buffer (50 mM Tris-HCl pH 7.4, 300 mM NaCl, 1 mM DTT, 2 mM MgCl_2_, 5% glycerol, 2 mM β-mercaptoethanol, 1 µl benzonase (Sigma), cOmplete EDTA-free protease inhibitors (Roche), CIP protease inhibitor (Sigma)). Cells were homogenized with a douncer. Cell lysates were cleared by centrifugation at 18,000 rpm for 45 min at 4°C in a SORVAL RC6+ centrifuge with an F21S-8×50Y rotor (Thermo Scientific). The supernatant was collected and incubated with pre-equilibrated Glutathione Sepharose 4B beads (GE Healthcare) for 2 h at 4°C with gentle shaking to bind GST-TEV-TBK1 or GST-TEV-EGFP-TBK1. Samples were centrifuged to pellet the beads and remove the unbound lysate. Beads were then washed five times with wash buffer (50 mM Tris-HCl pH 7.4, 300 mM NaCl, 5% glycerol, 1 mM DTT). Beads were incubated overnight with TEV protease in wash buffer (50 mM Tris-HCl pH 7.4, 300 mM NaCl, 5% glycerol, 1 mM DTT) at 4°C. After the proteins were released from the beads by the TEV protease, the supernatant was collected after centrifugation of the beads. The beads were washed twice with 4 ml of wash buffer, and the supernatant was collected. The supernatant fractions were pooled, filtered through a 0.45 µm syringe filter, and concentrated with a 30 kDa cut-off Amicon filter (Merck Millipore). The proteins were loaded onto a pre-equilibrated Superdex 200 Increase 10/300 GL column (Cytiva). Proteins were eluted with SEC buffer (25 mM Tris-HCl pH 7.4, 300 mM NaCl, 1 mM DTT). Fractions were analyzed by SDS-PAGE and Coomassie staining. Fractions containing purified TBK1 or GFP-TBK1 were pooled. After concentrating the purified protein, the protein was aliquoted and snap-frozen in liquid nitrogen. Proteins were stored at –80°C. A detailed protocol can be found here (https://doi.org/10.17504/protocols.io.81wgb6wy1lpk/v1).

To purify FIP200-GFP from insect cells, we purchased gene-synthesized codon-optimized GST-3C-FIP200-EGFP in a pGB-02-03 vector from Genscript (Addgene_187832). The V1 virus was generated as described above for SINTBAD. For expressions, we infected 1 L of Sf9 cells (12659017, Thermo Fisher, RRID:CVCL_0549), at 1 million cells per ml, with 1 ml of V1 virus. When the viability of the cells decreased to 90-95%, cells were collected by centrifugation. Cell pellets were washed with 1× PBS and flash-frozen in liquid nitrogen. Pellets were stored at –80°C. For purification of FIP200-GFP, the pellet was resuspended in 25 ml lysis buffer (50 mM HEPES pH 7.5, 300 mM NaCl, 1 mM MgCl_2_, 10% glycerol, 1mM DTT, 0.5% CHAPS, 1 µl benzonase (Sigma), cOmplete EDTA-free protease inhibitors (Roche), CIP protease inhibitor (Sigma)). Cells were homogenized with a douncer. Cell lysates were cleared by centrifugation at 72,000*g* for 45 min at 4°C with a Beckman Ti45 rotor. The supernatant was collected and incubated with pre-equilibrated Glutathione Sepharose 4B beads (GE Healthcare) for overnight at 4°C with gentle shaking to bind GST-3C-FIP200-EGFP. Samples were centrifuged to pellet the beads and remove the unbound lysate. Beads were washed seven times with wash buffer (50 mM HEPES pH 7.5, 200 mM NaCl, 1 mM MgCl_2_, 1 mM DTT). Beads were incubated overnight with precision 3C protease in wash buffer at 4°C. After the proteins were released from the beads by the 3C protease, the supernatant was collected after centrifugation of the beads. The beads were washed twice with 4 ml of wash buffer, and the supernatant was collected. The supernatant fractions were pooled, filtered through a 0.45 µm syringe filter, and concentrated with a 100 kDa cut-off Amicon filter (Merck Millipore). The proteins were loaded onto a pre-equilibrated Superose 6 Increase 10/300 GL column (Cytiva). Proteins were eluted with SEC buffer (25 mM HEPES pH 7.5, 200 mM NaCl, 1 mM DTT). Fractions were analyzed by SDS-PAGE and Coomassie staining. Fractions containing purified FIP200-GFP were pooled. After concentrating the purified protein, the protein was aliquoted and snap-frozen in liquid nitrogen. Proteins were stored at –80°C. A detailed protocol can be found here (https://doi.org/10.17504/protocols.io.dm6gpbkq5lzp/v1).

To purify the ULK1 complex (FIP200-ULK1-ATG13-ATG101) from HEK293 GnTI cells, we expressed and purified the complex in two parts. On one hand, we expressed the subcomplex FIP200-ATG13-ATG101 from pCAG vectors encoding GST-TEV-FIP200-MBP (RRID:Addgene_171410), ATG13 (RRID:Addgene_171412), GST-TEV-ATG101 (RRID:Addgene_171414). On the other hand, we expressed the ULK1 kinase from a pCAG backbone encoding MBP-TSF-TEV-ULK1 (RRID:Addgene_171416). The transfection and expression procedure was similar to what is described above for NAP1, with the exception that cells were harvested 48 h post-transfection. For purification of FIP200-ATG13-ATG101, the pellet was resuspended in 25 ml lysis buffer (50 mM Tris-HCl pH 7.4, 200 mM NaCl, 1 mM MgCl_2_, 10% glycerol, 1% Triton X-100, 1 mM TCEP, 1 µl benzonase (Sigma), cOmplete EDTA-free protease inhibitors (Roche), CIP protease inhibitor (Sigma)). Cells were homogenized with a douncer. Cell lysates were cleared by centrifugation at 18,000*g* for 30 min at 4°C with a SORVAL RC6+ centrifuge with an F21S-8×50Y rotor (Thermo Scientific). The supernatant was collected and incubated with pre-equilibrated Glutathione Sepharose 4B beads (GE Healthcare) for overnight at 4°C with gentle shaking to bind GST-TEV-FIP200-MBP. Samples were centrifuged to pellet the beads and remove the unbound lysate. Beads were washed seven times with wash buffer (50 mM HEPES pH 7.4, 500 mM NaCl, 1 mM MgCl_2_, 1 mM TCEP, 10% glycerol, 1% Triton X-100). Beads were incubated overnight with precision TEV protease in wash buffer at 4°C. After the proteins were released from the beads by the TEV protease, the supernatant was collected after centrifugation of the beads. The beads were washed twice with 4 ml of wash buffer, the supernatant was collected and pooled. For purification of MBP-TSF-TEV-ULK1, the cells were lysed and cleared as described for FIP200-ATG13-ATG101. The soluble supernatant was collected and incubated with pre-equilibrated Strep-Tactin Sepharose beads (IBA Life Sciences) for binding of the Twin-Strep-tagged ULK1 protein. After overnight incubation, the proteins were washed seven times with wash buffer (50 mM HEPES pH 7.5, 500 mM NaCl, 1 mM MgCl_2_, 1 mM TCEP, 10% glycerol, 1% Triton X-100). Beads were incubated overnight at 4°C with precision TEV protease in elution buffer (50 mM HEPES pH 7.4, 500 mM NaCl, 1 mM MgCl_2_, 1 mM TCEP, 10% glycerol, 1% Triton X-100). After the proteins were eluted from the beads, the supernatant was collected after centrifugation of the beads. The beads were washed twice with 4 ml of elution buffer, the supernatant was collected and pooled. The FIP200 and ULK1 samples were then subjected to a second step of affinity purification using the MBP tag. To this end, the FIP200-MBP-ATG13-ATG101 eluate and ULK1 eluate were mixed to allow reconstitution of the complex and loaded onto amylose resin (New England Biolabs). After 4 hours of incubation with the beads and extensive washing in washing buffer (20 mM HEPES pH 8, 200 mM NaCl, 2 mM MgCl_2_, 1 mM TCEP), proteins were eluted overnight at 4°C with gentle shaking in elution buffer (20 mM HEPES pH 8, 200 mM NaCl, 2 mM MgCl_2_, 1 mM TCEP, 50 mM Maltose). The final supernatant containing the purified protein complex was upconcentrated, aliquoted, and snap-frozen in liquid nitrogen. Proteins were stored at –80°C. Our protocol was based on this detailed description (https://doi.org/10.17504/protocols.io.bvn2n5ge).

### Microscopy-based bead assay

Glutathione Sepharose 4B beads (GE Healthcare) were used to bind GST-tagged bait proteins. To this end, 20 µl of beads were washed twice with dH_2_O and equilibrated with bead assay buffer (25 mM Tris-HCl pH 7.4, 150 mM NaCl, 1 mM DTT). Beads were then resuspended in 40 µl bead assay buffer, to which bait proteins were added at a final concentration of 5 µM. Beads were incubated with the bait proteins for 1 h at 4°C at a horizontal tube roller. Beads were then washed three times to remove unbound GST-tagged bait proteins and resuspended in 30 µl bead assay buffer. Where indicated, we also added MgCl_2_ and ATP to the buffer to allow the phosphorylation of targets by TBK1. Glass-bottom 384-well microplates (Greiner Bio-One) were prepared with 20 µl samples containing prey proteins at the concentrations described below and diluted in bead assay buffer, and 3 µl of beads were added per well. For the experiments in Figure 3D, NDP52 was used at a final concentration of 50 nM, FIP200-GFP, SINTBAD-mCherry, and TBK1 were used at a final concentration of 100 nM. For Figure 5B, mCherry-OPTN and GFP-TBK1 were used at a final concentration of 250 nM, and NAP1 was used from 100 nM to 10 µM. For Figure 6E, GFP-TBK1 was used at a final concentration of 250 nM. For Figure 7D, mCherry-OPTN, GFP-TBK1, and unlabeled NDP52 were used at a final concentration of 250 nM, while wild-type and mutant forms of NAP1 were used from 100 nM to 10 µM. For Figure S5, GFP-TBK1 was used at a final concentration of 250 nM. The beads were incubated with the prey proteins for 30 min prior to imaging, with the exception of Figure 3D, where proteins were co-incubated for 4 h before imaging. Samples were imaged with a Zeiss LSM 700 confocal microscope equipped with Plan Apochromat 20X/0.8 WD 0.55 mm objective. Three biological replicates were performed for each experimental condition.

For the quantification, we employed an artificial intelligence (AI) script that automatically quantifies signal intensities from microscopy images by drawing line profiles across beads and recording the difference between the minimum and maximum grey values along the lines. The AI was trained to recognize beads employing cellpose [80]. Processing is composed of two parts, with the first operating in batch mode. Multichannel input images are split into individual TIFF images and passed to cellpose (running in a Python environment). The labeled images produced by cellpose are re-assembled into multichannel images. Circular regions of interest (ROIs) are fitted to the segmented particles, and a pre-defined number of line profiles (here set to 20) are drawn automatically, starting at the center of the ROI and extending beyond the border of the circular ROI. This results in line profiles from the center of the bead into the inter-bead space of the well, allowing us to quantify the signal intensities at the rim of the beads. To prevent line profiles from protruding into adjacent beads, a combined ROI containing all beads was used. The AI-generated results were inspected manually for undetected beads, incorrect line profiles, or false-assigned bead structures. For each bead, a mean fluorescence and standard deviation are obtained based on the 20 line profiles per bead. Beads with standard deviations equal to or greater than half the mean value were either excluded or subjected to manual inspection for correction. To correct for inter-experiment variability in absolute values, the mean values for each bead were divided by the average bead intensity of the control condition. These values are then plotted and subjected to statistical significance calculations. A detailed protocol is available (https://doi.org/10.17504/protocols.io.14egn38pzl5d/v1).

### GFP pull down assay

GFP-tagged TBK1 was mixed with 20 µl of GFP-Trap agarose beads (Chromotek) at a final concentration of 1 µM. To this end, 20 µl of beads were washed twice with dH_2_O and equilibrated with bead assay buffer (25 mM Tris-HCl pH 7.4, 150 mM NaCl, 1 mM DTT). Beads were then resuspended in 40 µl bead assay buffer, to which GFP-TBK1 was added at a final concentration of 5 µM. Beads were incubated with GFP-TBK1 for 1 h at 4°C at a horizontal tube roller. Beads were washed three times to remove unbound GFP-tagged bait protein. Protein master mixes with prey protein were prepared in bead assay buffer at the following concentrations: mCherry-OPTN (1 µM), mCherry-NDP52 (1 µM), GST-NAP1 (1-10 µM). The protein master mixes were added to the beads and incubated for 1 h at 4°C at a horizontal tube roller. Beads were washed three times to remove unbound proteins, diluted in 60 µl of 1× Protein Loading dye, and heat-inactivated at 95°C for 5 min. Samples were analyzed by SDS-PAGE and Coomassie staining as described above. A detailed protocol is available (https://doi.org/10.17504/protocols.io.e6nvwd6x2lmk/v1).

### Quantification and statistical analysis

For the quantification of immunoblots, we performed a densitometric analysis using Fiji software. Graphs were plotted using Graphpad Prism version 9.5.1 (RRID:SCR_002798). For the quantification of microscopy-based bead assays, we employed an in-house developed AI tool to automate the recognition and quantification of the signal intensity for each bead, which resulted in a mean bead intensity value. These values were plotted and subjected to statistical testing. Depending on the number of samples, and as specified in the figure legends, we employed either a Student’s *t* test, a one-, or two-way ANOVA test with appropriate multiple comparison tests. Statistical significance is indicated with \**P*<0.05, \*\**P*<0.005, \*\*\**P*<0.001, \*\*\*\**P*<0.0001, ns, not significant. Error bars are reported as mean ± standard deviation. To ensure the reproducibility of experiments not quantified or subjected to statistical analysis, we showed one representative replicate in the paper of at least three replicates with similar outcomes for the main figures or at least two replicates for supplementary figures, as indicated in figure legends.

## Acknowledgments

We thank members of the Martens lab, Lazarou lab, James H. Hurley, Erika L. F. Holzbaur, Eunyong Park, Gerhard Hummer, Dorotea Fracchiolla, and other members of the Aligning Science Across Parkinson’s (ASAP) Team for their advice and helpful discussions. We thank the Max Perutz Labs BioOptics, Flow Cytometry, and Mass Spectrometry facilities for their technical support. We thank Jana Neuhold and the rest of the Vienna BioCenter Core Facilities (VBCF) Protech Facility for help with cloning and HEK cell expressions. We thank members from the Versteeg lab for training and assistance with lentiviral work. The summarizing schematic was generated with BioRender. This work was supported by a Marie Skłodowska-Curie MSCA Postdoctoral fellowship (101062916 to E.A.), a travel grant from the Flanders Fund for Scientific Research (FWO-Flanders to E.A.), the National Health and Medical Research Council (NHMRC) (GNT1106471 to M.L.), the Australian Research Council (ARC) Discovery Project (DP200100347 to M.L.). This research was also funded by Aligning Science Across Parkinson’s (ASAP). The Michael J. Fox Foundation for Parkinson’s Research (MJFF) administers the grant (ASAP-000350 to S.M. and M.L.) on behalf of ASAP and itself. For the purpose of open access, the authors have applied for a CC-BY public copyright license to the Author Accepted Manuscript (AAM) version arising from this submission.

## Author contributions

E.A., M.L., and S.M. conceived the project. E.A., T.N.N., B.S.P., M.L., S.M. designed the experiments. E.A., T.N.N., J.S.M., G.K., M.S., S.S., E.M.W., K.D.C., and B.S.P. performed the experiments. E.A. and S.M. wrote the original draft to which all authors contributed by editing and reviewing.

## Author ORCID IDs

Elias Adriaenssens (0000-0001-9430-917X)

Thanh Ngoc Nguyen (0000-0001-9698-0020)

Justyna Sawa-Makarska (0000-0002-9321-976X)

Grace Khuu (0000-0002-2550-0605)

Martina Schuschnig (not available)

Emily Maria Watts (not available)

Stephen Shoebridge (0000-0002-3555-2855)

Kitti Dora Csalyi (0000-0002-0853-8257)

Benjamin Scott Padman (0000-0002-5710-6100)

Michael Lazarou (0000-0003-2150-5545)

Sascha Martens (0000-0003-3786-8199)

## Data availability statement

Raw files associated with this work will be made available on Zenodo by time of final publication.

## Code availability statement

### Supplementary figures

**Figure S1.**
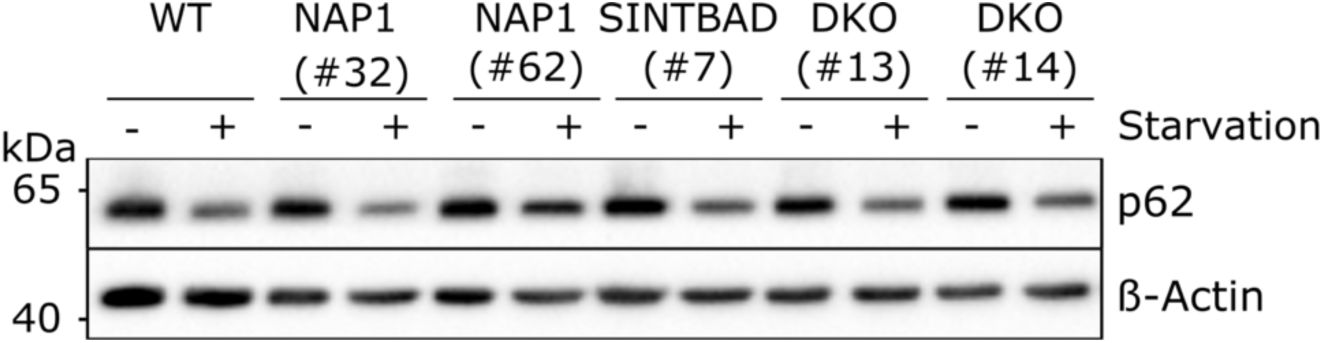
NAP1 and SINTBAD are not essential for non-selective starvation autophagy. Immunoblotting of p62 levels in wild-type (WT), NAP1 knockout, SINTBAD knockout, and NAP1/SINTBAD double knockout (DKO) HeLa cells expressing YFP-Parkin, untreated or treated with EBSS starvation medium for 8 h.

**Figure S2.**
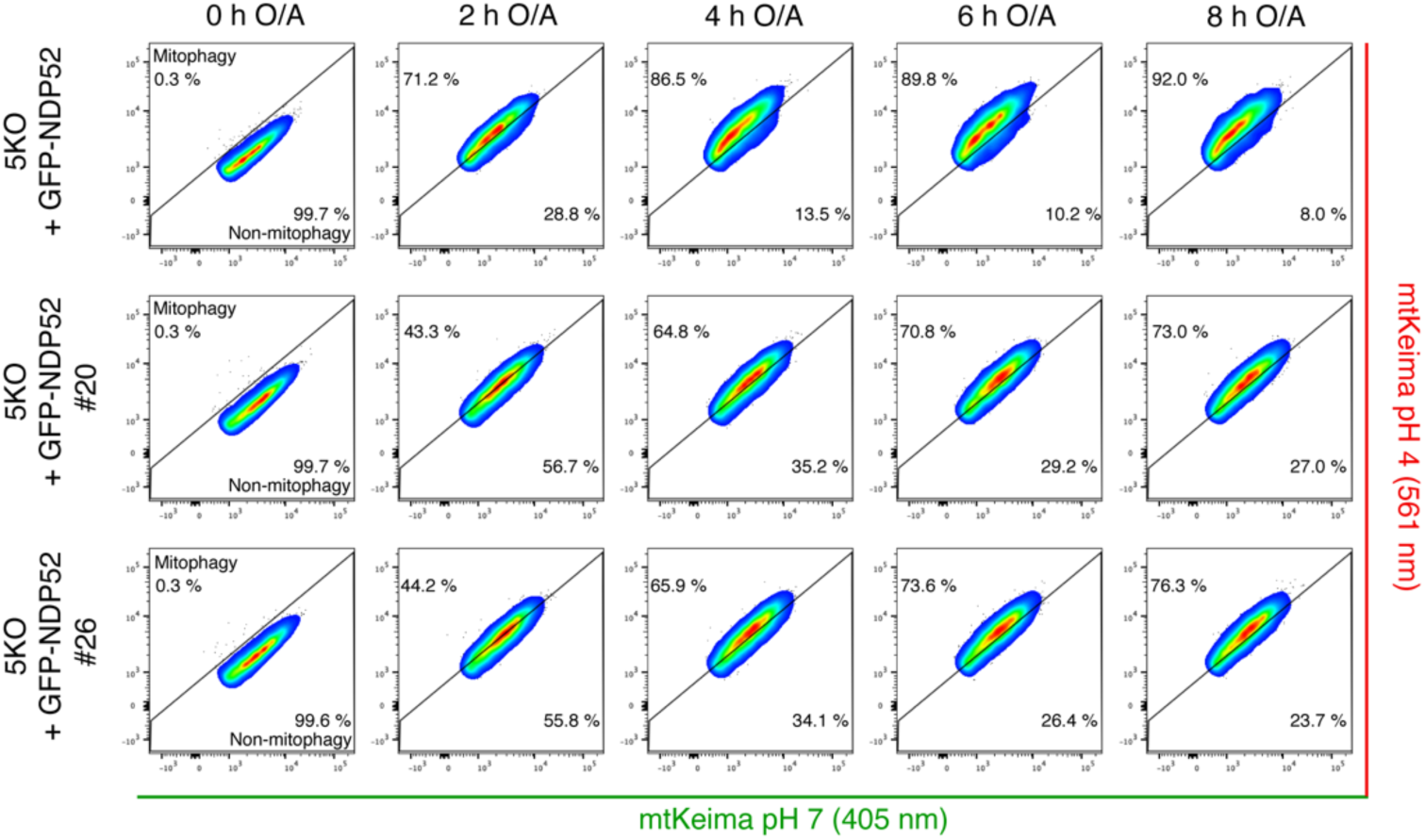
Mt-mKeima mitophagy flux assay in pentaKO cells rescued with NDP52 and in the presence or absence of NAP1/SINTBAD. Representative replicate showing reduced mitophagy initiation in both NAP1/SINTBAD DKO clones (#20 and #26) in a pentaKO (5KO) background, expressing BFP-Parkin, and rescued with GFP-NDP52. Cells were either untreated (time point 0 h) or treated with O/A for the indicated times. The mt-mKeima signal was analyzed by flow cytometry and quantified. Results are representative of one of three replicates.

**Figure S3.**
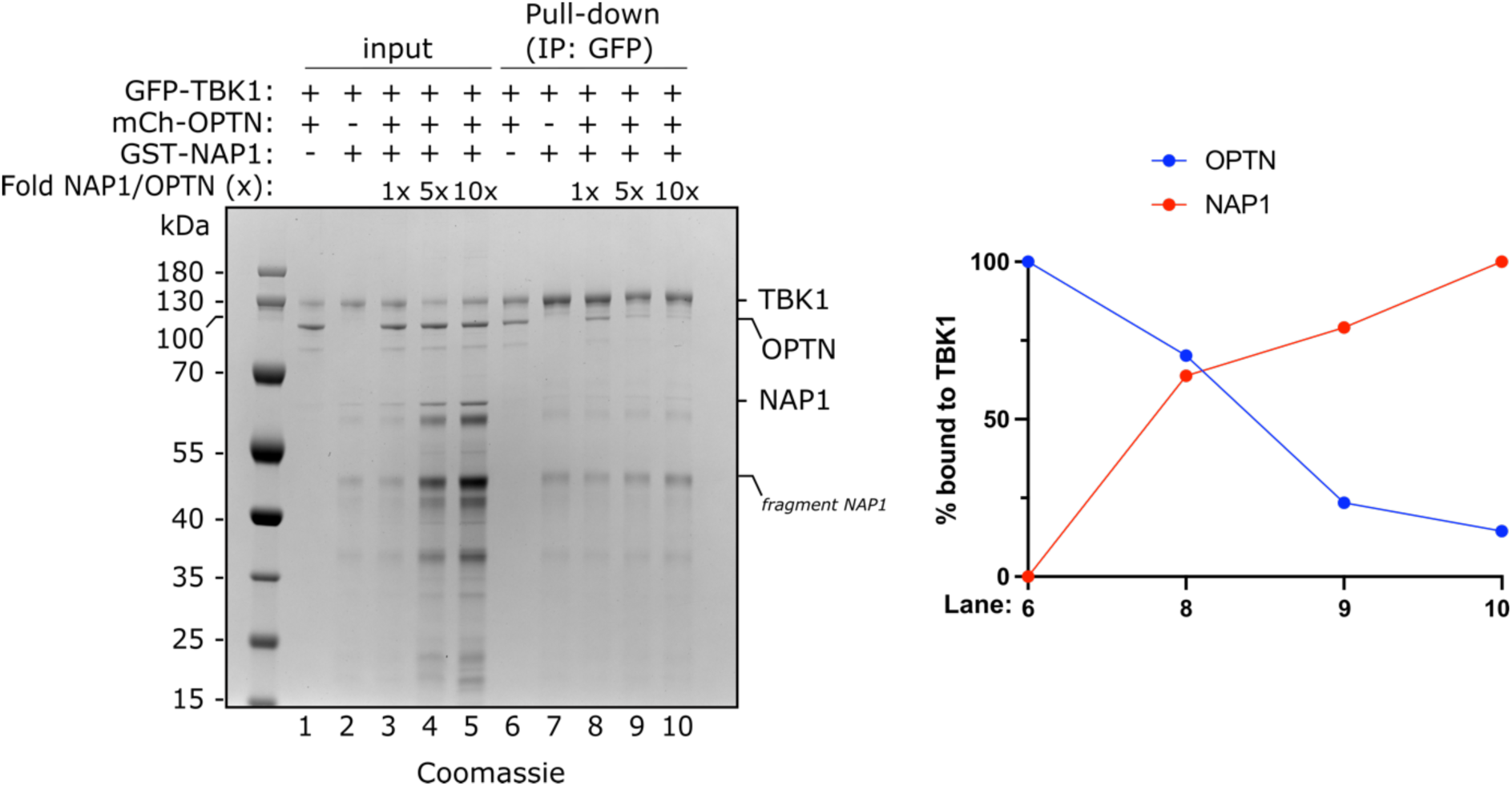
Pull-down of GFP-TBK1 with mCherry-OPTN and GST-NAP1. Pull-down assay of mCherry-OPTN versus GST-NAP1 by GFP-TBK1. GFP-TBK1 was pre-loaded onto GFP-Trap beads and then incubated with the protein mixtures as indicated. The relative amounts of mCherry-OPTN and GST-NAP1 bound to TBK1 were quantified for the indicated lanes and plotted (right).

**Figure S4.**
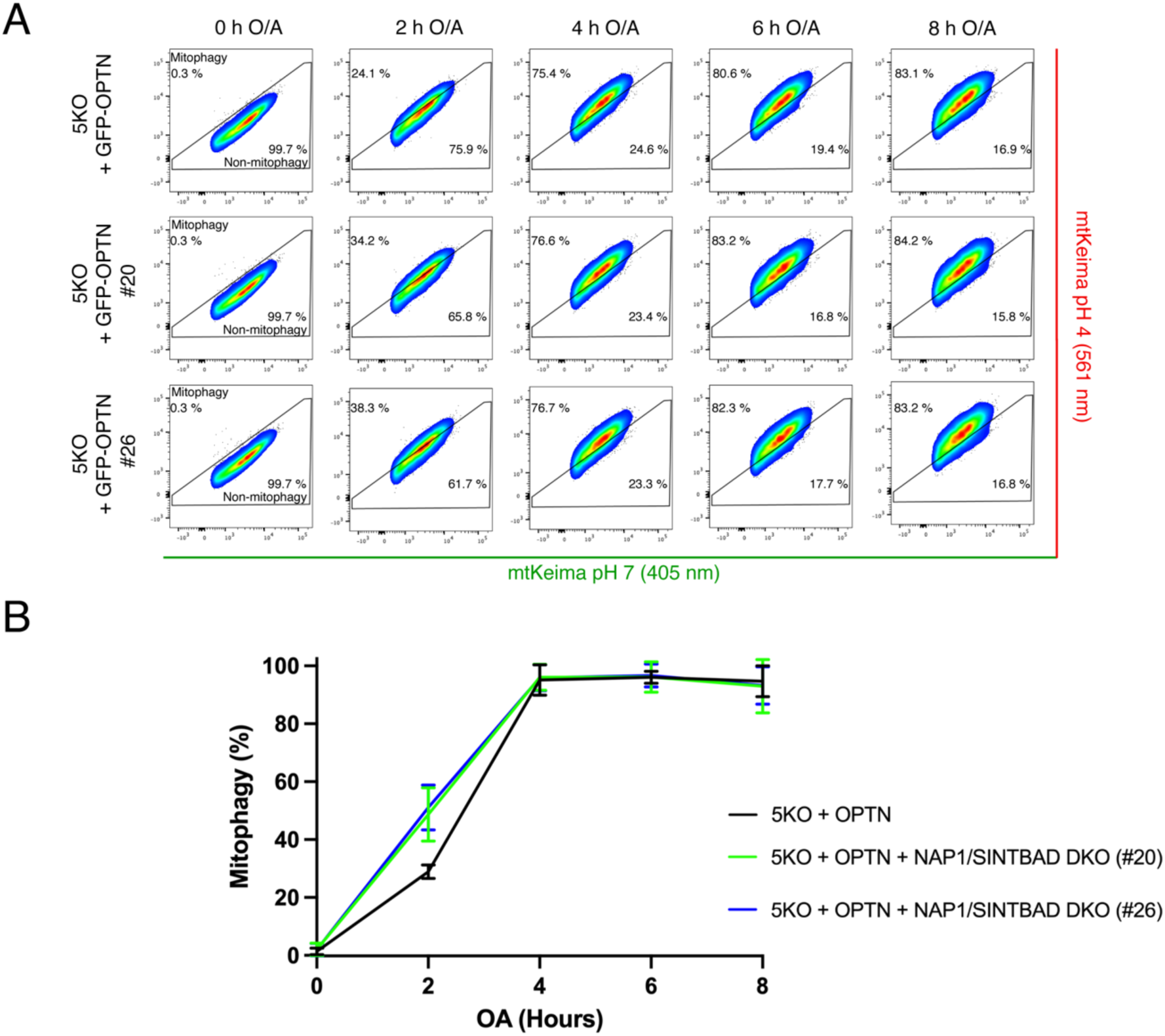
Mt-mKeima mitophagy flux assay in pentaKO cells rescued with OPTN and in presence or absence of NAP1/SINTBAD. (**A-B**) pentaKO (5KO) and NAP1/SINTBAD DKO/5KO (clones #20 and #26)) expressing BFP-Parkin, GFP-OPTN and mt-mKeima were left untreated and treated with O/A for indicated time points and mitophagy flux was measured via FACS (A) and quantified (B) (n=3).

**Figure S5.**
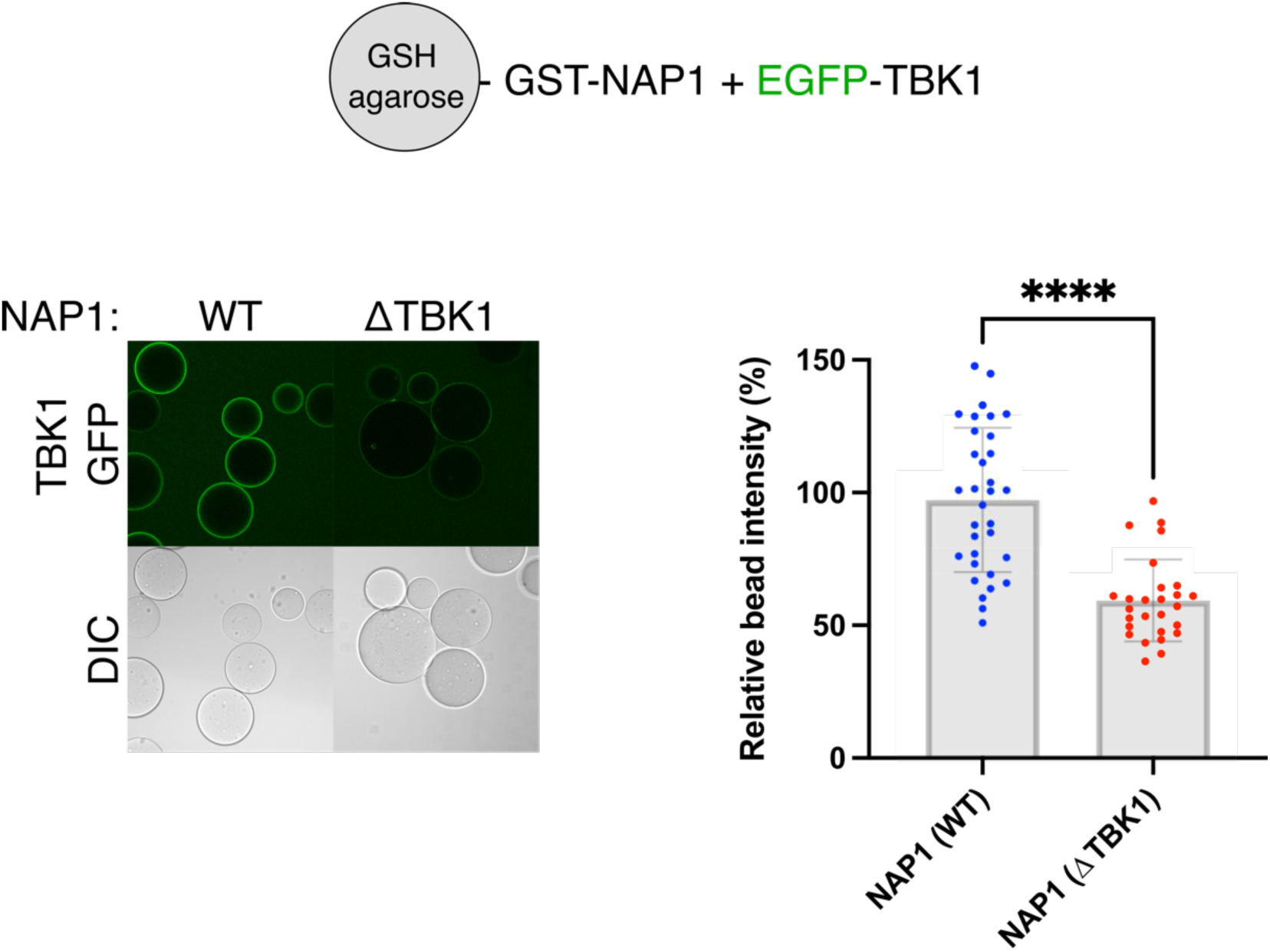
*In vitro* TBK1 binding assay of NAP1 (L226Q/L233Q) mutant. In vitro binding assay using glutathione-coupled agarose beads coated with GST-NAP1 wild-type (WT) or the ΔTBK1 (L226Q/L233Q) mutant and incubated with EGFP-TBK1. Samples were analyzed by confocal microscopy. Data in are shown as mean ± s.d. from three independent experiments. Each data point represents the mean signal intensity for an individual bead. A *t* test was performed. \*\*\*\**P*<0.0001.

**Figure S6.**
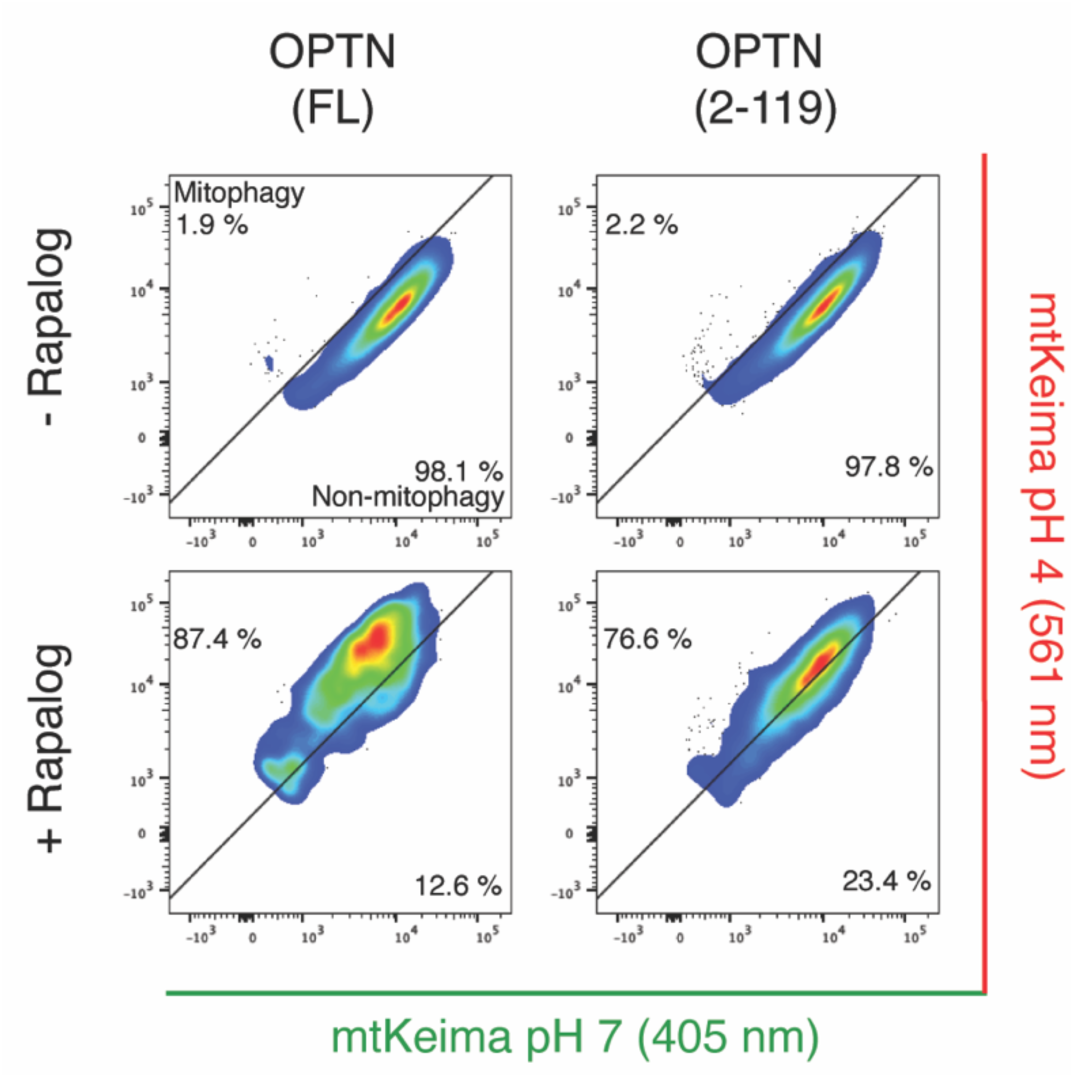
Chemical-dimerization assay with full-length OPTN and OPTN (2-119) PentaKO HeLa cells expressing mt-mKeima, Fis1-FRB and FKBP-OPTN wild-type (WT) or amino acids 2-119 (2-119) were left untreated or treated with rapalog for 24 hours as indicated. The mitophagy flux was analyzed by flow cytometry.

**Figure S7.**
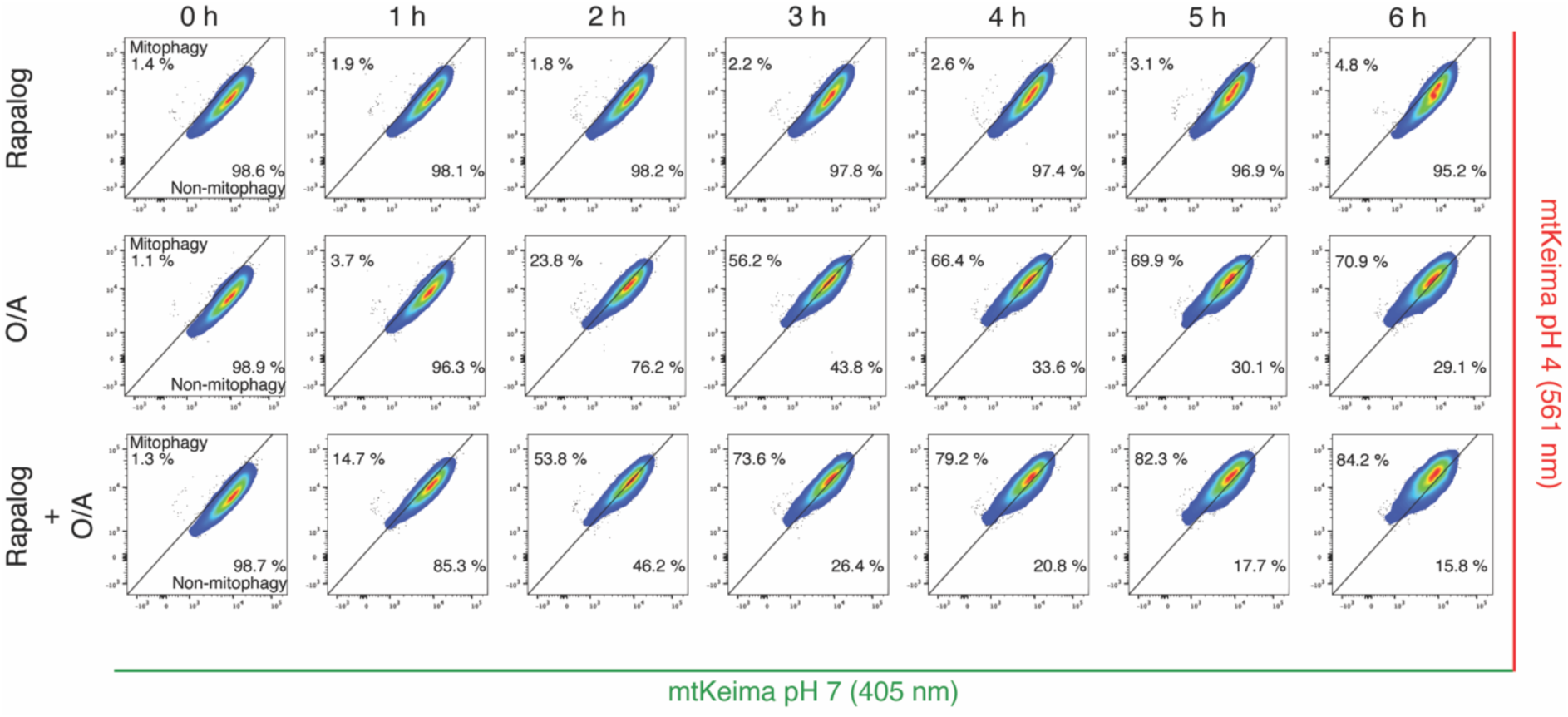
Chemical-dimerization assay with OPTN (2-119) in pentaKO HeLa cells rescued with GFP-NDP52. PentaKO HeLa cells expressing BFP-Parkin, GFP-NDP52, and mt-mKeima were further transduced with Fis1-FRB and FKBP-OPTN amino acids 2-119 (2-119). Cells were either left untreated (time point 0 h) or treated with rapalog alone, O/A alone, or rapalog plus O/A for the indicated times. The mt-mKeima signal was measured by flow cytometry.

